# Extensive transfer of genes for edible seaweed digestion from marine to human gut bacteria

**DOI:** 10.1101/2020.06.09.142968

**Authors:** Nicholas A. Pudlo, Gabriel Vasconcelos Pereira, Jaagni Parnami, Melissa Cid, Stephanie Markert, Jeffrey P. Tingley, Frank Unfried, Ahmed Ali, Austin Campbell, Karthik Urs, Yao Xiao, Ryan Adams, Duña Martin, David N. Bolam, Dörte Becher, Thomas M. Schmidt, D. Wade Abbott, Thomas Schweder, Jan Hendrik Hehemann, Eric C. Martens

**Affiliations:** Department of Microbiology and Immunology, University of Michigan, Ann Arbor, MI 48109, USA; Max Planck Institute for Marine Biology, Bremen, Germany; University of Bremen, Center for Marine Environmental Sciences (MARUM), 28359 Bremen, Germany; Pharmaceutical Biotechnology, University of Greifswald, 17487 Greifswald, Germany; Institute of Marine Biotechnology, 17489 Greifswald, Germany; Agriculture and Agri-Food Canada, Lethbridge Research and Development Centre, Lethbridge, AB, Canada; Institute for Cell and Molecular Biosciences, Newcastle University, Newcastle upon Tyne, UK; Department of Internal Medicine, University of Michigan, Ann Arbor, MI 48109, USA; Institute of Microbiology, University Greifswald, 17487 Greifswald, Germany

## Abstract

Humans harbor numerous species of colonic bacteria that digest the fiber polysaccharides in commonly consumed terrestrial plants. More recently in history, regional populations have consumed edible macroalgae seaweeds containing unique polysaccharides. It remains unclear how extensively gut bacteria have adapted to digest these nutrients and use these abilities to colonize microbiomes around the world, especially outside Asia. Here, we show that the ability of gut bacteria to digest seaweed polysaccharides is more pervasive than previously appreciated. Using culture-based approaches, we show that known *Bacteroides* genes involved in seaweed degradation have mobilized into many members of this genus. We also identify several previously unknown examples of marine bacteria-derived genes, and their corresponding mobile DNA elements, that are involved in degrading seaweed polysaccharides. Some of these genes reside in gut-resident, Gram-positive Firmicutes, for which phylogenetic analysis suggests an origin in the *Epulopiscium* gut symbionts of marine fishes. Our results are important for understanding the metabolic plasticity of the human gut microbiome, the global exchange of genes in the context of dietary selective pressures and identifying new functions that can be introduced or engineered to design and fill orthogonal niches for a future generation of engineered probiotics.

---

More than any other animal on Earth, as humans have extended into new territory and developed civilization, we have expanded our nutritional repertoire out of either necessity, curiosity or the pleasure of cuisine. One example is the consumption of edible red and brown macroalgae seaweeds, originally by populations living near coasts, which introduced a variety of novel dietary fiber polysaccharides such as laminarin, alginate, carrageenan, agarose and porphyran (**Fig. 1A**). Compared to commonly consumed fruits, vegetables, grains and nuts, the polysaccharides in seaweeds have different chemical structures that require specific degradative enzymes^1,2^. As with nearly all dietary fibers, humans lack these enzymes and therefore rely on colonic bacteria for the ability to digest them. Previous studies have documented a few examples in which symbiotic bacteria belonging to *Bacteroides—*a dominant genus in humans—possess genes for degrading seaweed-derived porphyran^3,4^, agarose^5^, alginate^6,7^ and laminarin^8^. In the first three cases, the genes involved often have closest homologs in marine Bacteroidetes, which are physiologically different from gut *Bacteroides* but share similar mechanisms for polysaccharide degradation^9^. In the case of porphyran degradation, a mobile DNA element was identified that presumably carried the required genes into the human microbiome, integrating in a site-specific manner into the genomes of at least two species^3,4,10^. However, it remains unknown how pervasively genes involved in seaweed degradation have permeated the human gut microbiome, the sources and taxonomic range of these “genetic upgrades” from environmental bacteria and the DNA transfer mechanisms involved.

**Figure 1.**
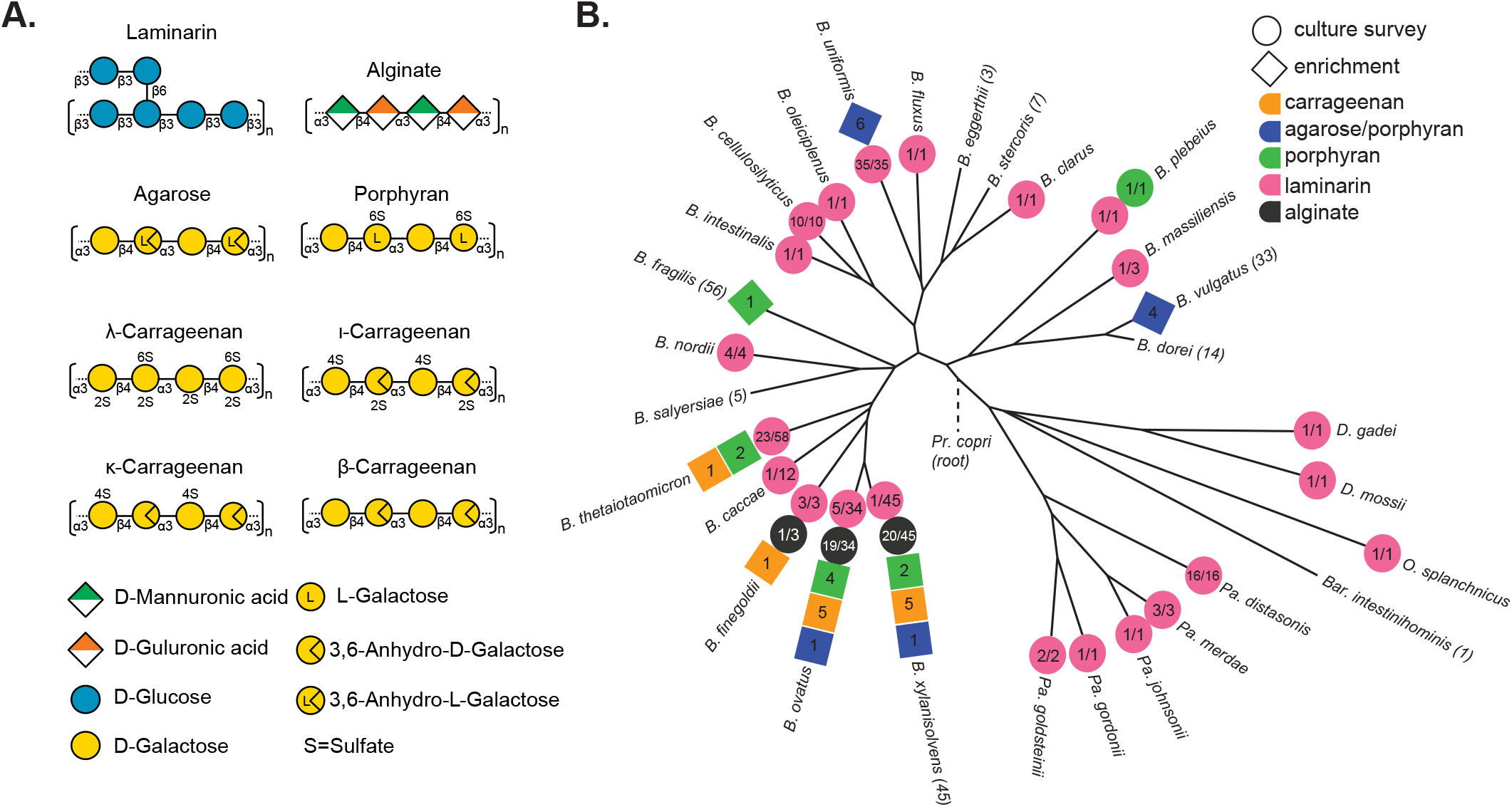
(**A**) Structures of the 5 different seaweed-derived polysaccharides used in this study. (**B**) A phylogeny of human gut Bacteroidetes based on core gene alignment^60^ and illustration of seaweed polysaccharide-degrading abilities present in members of each species. Degradative abilities that were observed in isolates cultured without targeted enrichment are represented as circles and the numbers indicate the number of positive strains over the total number tested. Isolates recovered by enrichment on agarose, porphyran or carrageenan are shown in diamonds and the number indicates the total number of isolates recovered for that species from different donors.

## Distribution of algal polysaccharide degradation in human gut Bacteroidetes

As a first step to determine the extent to which gut bacteria have acquired seaweed degrading abilities, we surveyed a collection of human and animal gut Bacteroidetes representing 30 different species (354 total isolates, **Supplemental Table 1**) on four of the five algal polysaccharides mentioned above (all except agarose due to solubility problems in high-throughput). The ability to use the β-linked glucan laminarin from *Laminaria digitata*, for which we recently identified the corresponding genes in strains of *Bacteroides uniformis* and *Bacteroides thetaiotaomicron*^8^, was broadly represented and present in members of 22 different species (**Fig. 1B**). This may reflect the ubiquity of structurally related, but non seaweed-derived, β-linked glucans in many different plants and fungi, albeit with subtle variations in structure^8^.

The remaining seaweed polysaccharides were used by far fewer species. Alginate was the second most prevalent trait but was confined to a subset of strains belonging to just three related species (**Fig. 1B)**. Among strains that grew on alginate, all for which a sequenced genome was available contained known alginate utilization genes^6,7^ and loci from representatives in each species were upregulated >100-fold in response to growth on alginate (**Extended Data 1**). Growth on the remaining polysaccharides was even less prevalent, with only the single, previously known isolate of *Bacteroides plebeius* from a Japanese adult showing growth on porphyran and individual isolates belonging to two different species, *B. thetaiotaomicron* and *B. ovatus*, growing on carrageenan (a mixture of κ, λ isomers).

## Carrageenan utilization is associated with mobile DNA

Unlike agarose and porphyran, which have been traditionally consumed in Asian cuisine, the red algae that contain carrageenan (CGN) have been associated with both European and Chinese consumption as far back at 400 BC and CGN is now a widely used food additive^11^. To determine the genes involved in CGN utilization—an ability that has not yet been described in gut bacteria—we grew *B. thetaiotaomicron* strain 3731 (herein, *Bt*^3731^) on λ-CGN and performed RNA-seq-based transcriptional profiling and proteomics to identify upregulated functions compared to growth on galactose. A total of 343 genes were upregulated based on the criteria used (**Supplemental Table 2**) and 56 of these genes were contained in two fragments of a polysaccharide utilization locus (PUL), a hallmark of Bacteroidetes nutrient assimilation (**Fig. 2A**)^2^. More than half of these genes had nearest homologs in Gram-negative marine bacteria, suggesting transfer from marine species that harbor genes for CGN utilization (**Supplemental Table 3**)^12^. Concurrent proteomics analysis revealed strong concordance with the transcriptome, demonstrating that *Bt*^3731^ dedicates substantial resources to production of 44 different proteins that are increased greater than 0.01% of normalized spectral abundance factor (NSAF) in CGN (**Fig. 2D, Supplemental Table 4**). Finally, a similar PUL was identified in the genome of CGN-degrading *B. ovatus* CL02T12C04 (herein, *Bo*^12C04^) the second strain in our survey that grew on this seaweed polysaccharide (**Fig. 2B, Extended Data 2**).

**Figure 2.**
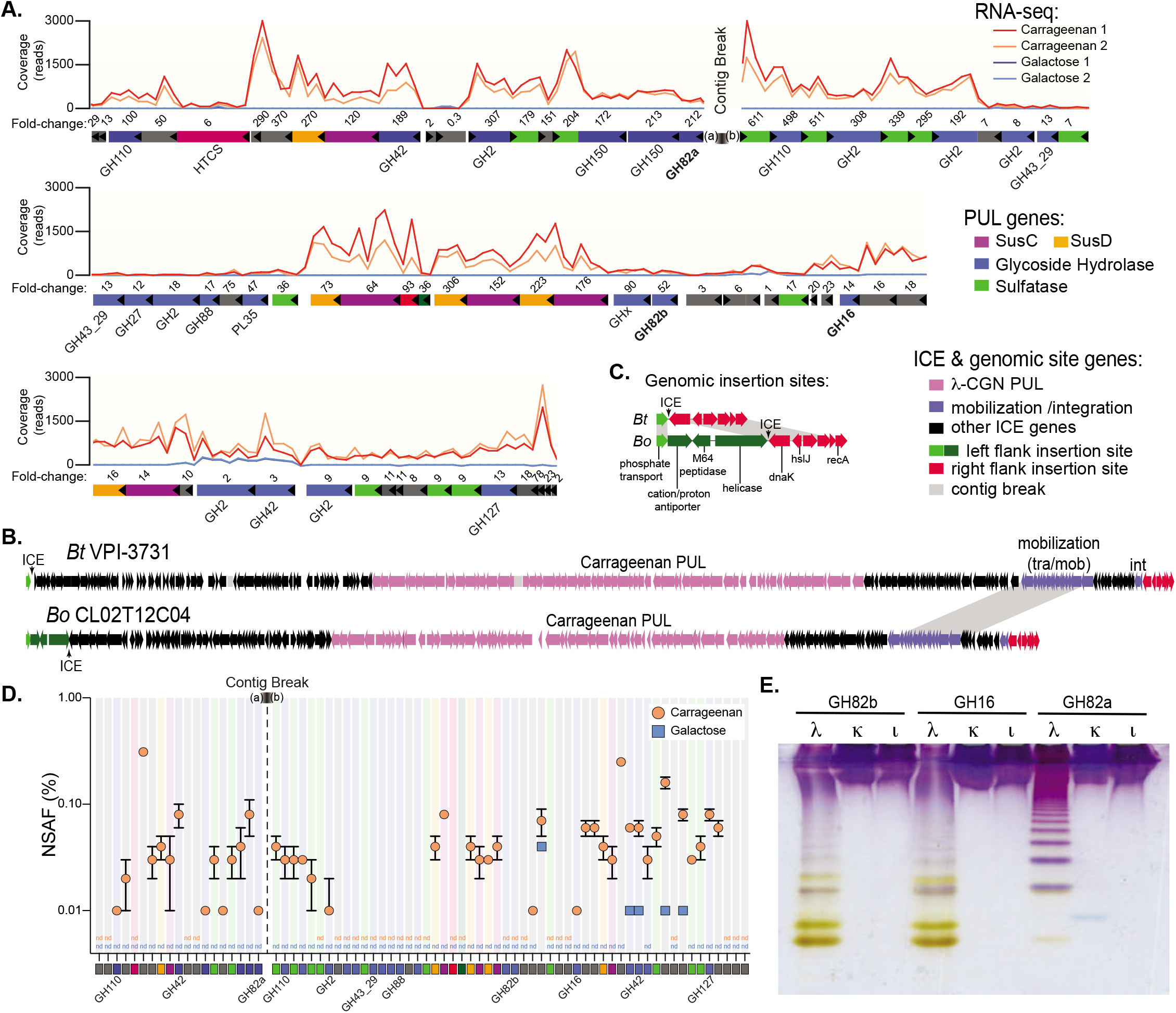
**(A)** Transcriptional responses of genes within the 130.5kbp PUL in *Bt*^3731^ during growth on λ-CGN compared to a galactose reference. **(B)** A comparison of the larger *Bt*^3731^ and *Bo*^12C04^ ICE gene architecture showing the flanking chromosomal backbone genes in green and red. **(C)** Higher resolution view of the *Bt*^3731^ and *Bo*^12C04^ ICE insertion sites near the conserved *dnaK* gene. **(D)** Relative abundances (in % of normalized spectral abundance factor – NSAF) of PUL-encoded proteins detected in the total protein fraction of cultures grown on carrageenan (orange) and galactose (blue). Error bars are the standard error of the mean (SEM). Proteins that could be detected in at least two out of three independent biological replicates of each substrate condition are shown. **(E**) C-PAGE gel showing digestion of three different CGN isomer types with recombinant GH16 and GH82 enzymes that are highly expressed within the *Bt*^3731^ PUL. Shorter CGN oligosaccharides migrate toward the bottom of the gel.

The genes immediately adjacent to the newly identified CGN PULs are homologous in *Bt*^3731^ and *Bo*^12C04^ and suggest lateral gene transfer. In particular, we observed a contiguous set of mobilization genes near one side of the PUL (**Fig. 2B**), indicating that this region is an integrative conjugative element (ICE). To test this further, we performed comparative genomics with several *B. thetaiotaomicron* and *B. ovatus* strains that do not degrade CGN, revealing the presence of identical 16 bp direct repeat sequences at the ends of each putative ICE. Both flanking genomic regions were syntenic between CGN-degrading and non-degrading isolates with only a single copy of the 16 bp repeat in non-degraders that was immediately downstream of the housekeeping gene *dnaK* (**Fig. 2C**). PCR and sequence analysis of this region demonstrated the ability of this element to circularize from the *Bt*^3731^ genome (**Extended Data 1**), providing support for the conclusion that *Bacteroides* carrageenan utilization was conferred by acquisition of a large (245-265 kbp) ICE.

Analysis of recombinant *Bt*^3731^ enzymes from the putative CGN PUL supported its role in degrading this seaweed polysaccharide. A recombinant glycoside hydrolase family 16 (GH16) enzyme, and two GH82 enzymes, revealed that they are each *endo*-acting carrageenases with specificity for λ-CGN (**Fig. 2E**), which is consistent with additional growth data for this strain and members of 3 other species described below. Interestingly, *Bt*^3731^ that were actively growing on λ-CGN released oligosaccharides in a similar size range as those produced by recombinant *endo*-acting enzymes (**Extended Data 2**). This is notable because degraded forms of CGN, so called “poligeenan”, have been associated with intestinal inflammation and colorectal cancer development in animal models^13^, leading to the exclusion of low molecular weight dietary CGN in many countries^14^. Thus, it is possible that some humans harbor CGN-degrading *Bacteroides* that might convert the non-harmful high molecular weight form of this polysaccharide into harmful oligosaccharides during catabolism, a metabolic transformation that requires additional investigation.

## Enrichment culture to isolate new seaweed degrading strains

The presence of porphyran and CGN utilization was rare (0.3-0.6%) in our collection of human and animal gut Bacteroidetes. This observation could be explained by geographic bias in the humans from which they were cultured or latent populations of bacteria that are enriched only after seaweed consumption. As an additional approach to identify bacteria that possess these traits, we surveyed stool samples from 240 different healthy volunteers using an enrichment strategy (see *Methods*) that was previously successful in isolating an agarose-degrading strain of *B. uniformis*^3,5^. Using this approach, we isolated 33 new strains that were enriched based on the ability to grow on CGN, porphyran or agarose (**Extended Data 3**). Identification by 16S rDNA gene sequencing revealed that these isolates belong to 6 different species (**Fig. 1B**). CGN utilization was confined to members of four related species, including the two identified above and extending to *B. xylanisolvens* and *B. finegoldii*. Consistent with the recombinant enzyme data shown above, growth on CGN was specific for λ-CGN and not the i or k isomers (**Extended Data 3**), which vary in sulfate position (**Fig. 1A**). All of the isolates that grew on agarose were also capable of some growth on the porphyran preparation used, which is likely since porphyran is a heteropolysaccharide that contains agarose segments with 3,6 anhydro-L-galactose instead of L6-S-galactose found in the material purified from *Porphyra yezoensis^10^*. However, since porphyran growth was sometimes better than agarose in these strains (*e.g*., *B. xylanisolvens* AO201-2, **Extended Data 3**) it is possible that the sensory and/or enzymatic machinery in some of these strains is optimized towards porphyran. Such combined agarose/porphyran utilization was previously observed in *B. uniformis* ^5^ and also observed here in three more species, *B. vulgatus, B. ovatus* and *B. xylanisolvens*. Utilization of only porphyran was broadly distributed across the *Bacteroides* genus and, including previously identified isolates^4,10^, is now known to be present in six different species.

To determine if the new isolates harbor genes for seaweed utilization that are related to the previously identified PULs and associated mobile elements, we performed draft genome sequencing on each isolate and assembled the reads using the known gene clusters as scaffolds. All of the newly identified CGN-degrading strains had regions with high coverage (62-100% of PUL length) and nearly identical homology (>99% nt identity) when mapped to the known *Bo*^12C04^ PUL, the closest match for all new isolates (**Extended Data 4)**. This suggests that the new CGN degrading strains all contain close variants of the genes identified above.

We previously used phylogenetic analysis of a common *Bacteroides* conjugation protein (TraJ) to identify groups of ICEs that are associated with various genetic cargo (including PULs), examine their relationship to one another and identify integration site preferences^15^. This analysis revealed that the original *B. plebeius* ICE that confers porphyran utilization and integrates into tRNA^Lys(TTT)^ is closely related in both sequence and architecture to a family of ICEs that integrate into tRNA^Phe(GAA)^ and carry variable gene cargo, including a *B. thetaiotaomicron* PUL for fungal α-mannan degradation^16^. To determine if the transfer functions associated with *Bacteroides* CGN ICEs are related to the previously characterized ICEs, we created a new phylogeny of 2,255 TraJ sequences from currently sequenced gut, marine and environmental Bacteroidetes (**Extended Data 4**). The TraJ proteins associated with the CGN ICEs are only distantly related to those involved in porphyran transfer to *B. plebeius*, sharing 22% amino acid identity and supporting the conclusion that distinct mobile elements have captured their respective genetic cargo for degrading these seaweed polysaccharides.

All of the new agarose/porphyran utilizing strains contained sequences that were nearly identical (>99% nt homology) to the *B. uniformis* PUL that was previously demonstrated to be involved predominantly in agarose degradation (**Extended Data 5**)^5^. However, the surrounding genome region was not resolved for this PUL in the initial strain that was sequenced. Interestingly, a recent *B. uniformis* isolate that was recovered from a healthy Japanese adult and sequenced using long-read sequencing, also contains this known agarose PUL, which was resolved as part of a 124 kbp circular plasmid. Using this plasmid template, we mapped sequences from all 11 new strains, finding that each contains sequences that match to between 46.2-94.2 % of this plasmid, with all of the *B. uniformis* and one *B. xylanisolvens* strain containing most of the episome (**Fig. 3**). Interestingly, a previously identified plasmid containing agarolytic genes was isolated from a marine sediment Bacteroidetes strain but found to be non-conjugal and lack mobilization machinery^17^. A potentially self-replicating plasmid that transfers polysaccharide utilization functions among Bacteroidetes has not yet been described and raises new possibilities for biotechnological applications. The mobilization genes that were contained on this plasmid were also divergent from the examples discussed above and lack TraJ homologs and could not be placed in our phylogeny. However, they underscore the diversity of mechanisms that have been coopted to capture useful genetic cargo and transfer it between related bacteria that exist in different environments.

**Figure 3.**
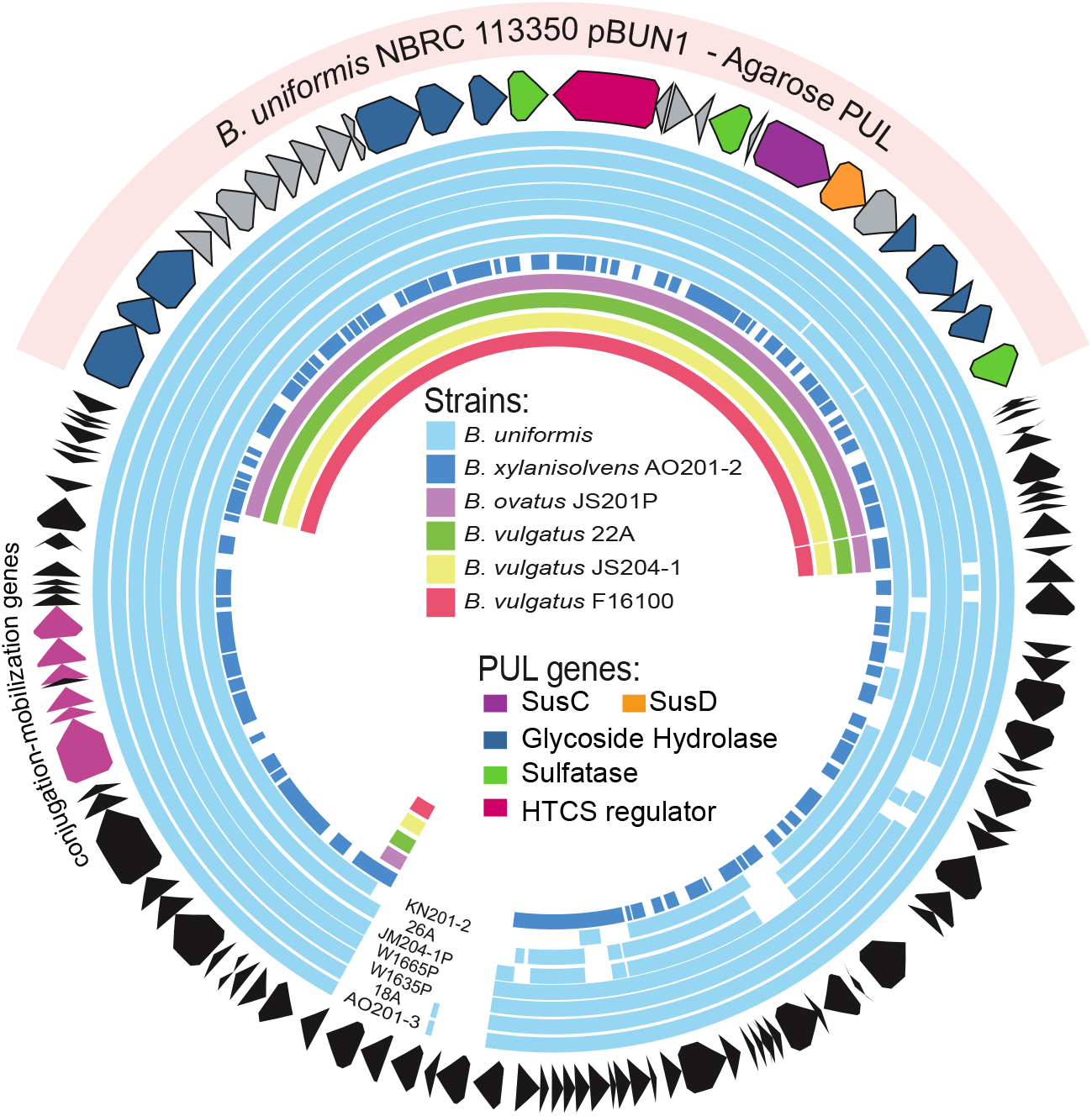
Circular map of the *B. uniformis* strain NBRC 113350 plasmid pBUN1 that contains an agarose utilization PUL. Sequence reads from each of the additional isolates in this study were mapped based on a minimum of 97% nucleotide identity to this scaffold.

## A second lateral transfer event for porphyran utilization into *Bacteroides*

All but one of the identified porphyran-degrading strains had sequences with high coverage (75-100%) and nucleotide identity (>96%) to the known ICE originally identified in *B. plebeius^3^* (**Extended Data 5**). Thus, variants of this mobile DNA have successfully integrated into at least 6 different species. Interestingly, one newly isolated *B. xylanisolvens* strain (*Bx*^18-002P^) did not contain extensive sequence homology to the previously identified porphyran ICE (**Fig. 4A**), suggesting that its growth on porphyran is attributable to an alternative set of genes. To explore this, we generated a high-quality, nearly closed genome for this isolate and performed RNA-seq on cells grown on porphyran compared to a galactose reference. A notably large set of 96 genes within two closely linked chromosomal regions showed strong responses (**Supplemental Table 5**). These genes also have close relatives in marine Bacteroidetes (**Supplemental Table 6**) and encode *Bacteroides* PUL functions, including enzymes predicted to degrade porphyran, such as families GH16, GH86 and GH117, sulfatases and many other enzymes (**Fig. 4B**). Comparison of these gene clusters to another recently cultured and sequenced *B. xylanisolvens* (*Bx*^BIOML-A58^) revealed that this second strain—which was presumably not isolated based on its ability to grow on porphyran^18^—also contained the same genes that were porphyran-inducible in *Bx*^18-002P^. However, this strain lacked a region present in *Bx*^18-002P^ that separates the two blocks of porphyran-responsive genes and contains several mobilization genes. In addition, both strains have syntenic and homologous genes on both sides of the porphyran-inducible PUL, with additional mobilization genes contained on one side (**Fig. 4D**). Based on this, we are unable to conclude which, if any, of the mobilization genes might be associated with carrying these newly identified porphyran-degrading functions into *Bx*^18-002P^ and *Bx*^bIOML-A58^. However, the simplest scenario is that the common DNA shared between these strains represents an ancestral ICE architecture and strain *Bx*^18-002P^ harbors a second, unrelated ICE that integrated into the middle of a large porphyran PUL without disrupting its function. As with the CGN-degrading strains, comparison to other *Bx* strains that lack the ability to grow on porphyran suggested that an ancestral genomic region, in this case encoding a nearby tRNA^Lys(TTT)^ and thymidine kinase (*tdk*) gene was the site of ICE integration via a 21 bp direct repeat, which interestingly targets the same tRNA as the first porphyran ICE but via a different sequence (**Fig. 4C**). This idea was supported by PCR and sequence analysis on these regions, which revealed that this ICE is capable of excising to restore the predicted ancestral genomic architecture (**Extended Data 1**).

**Figure 4.**
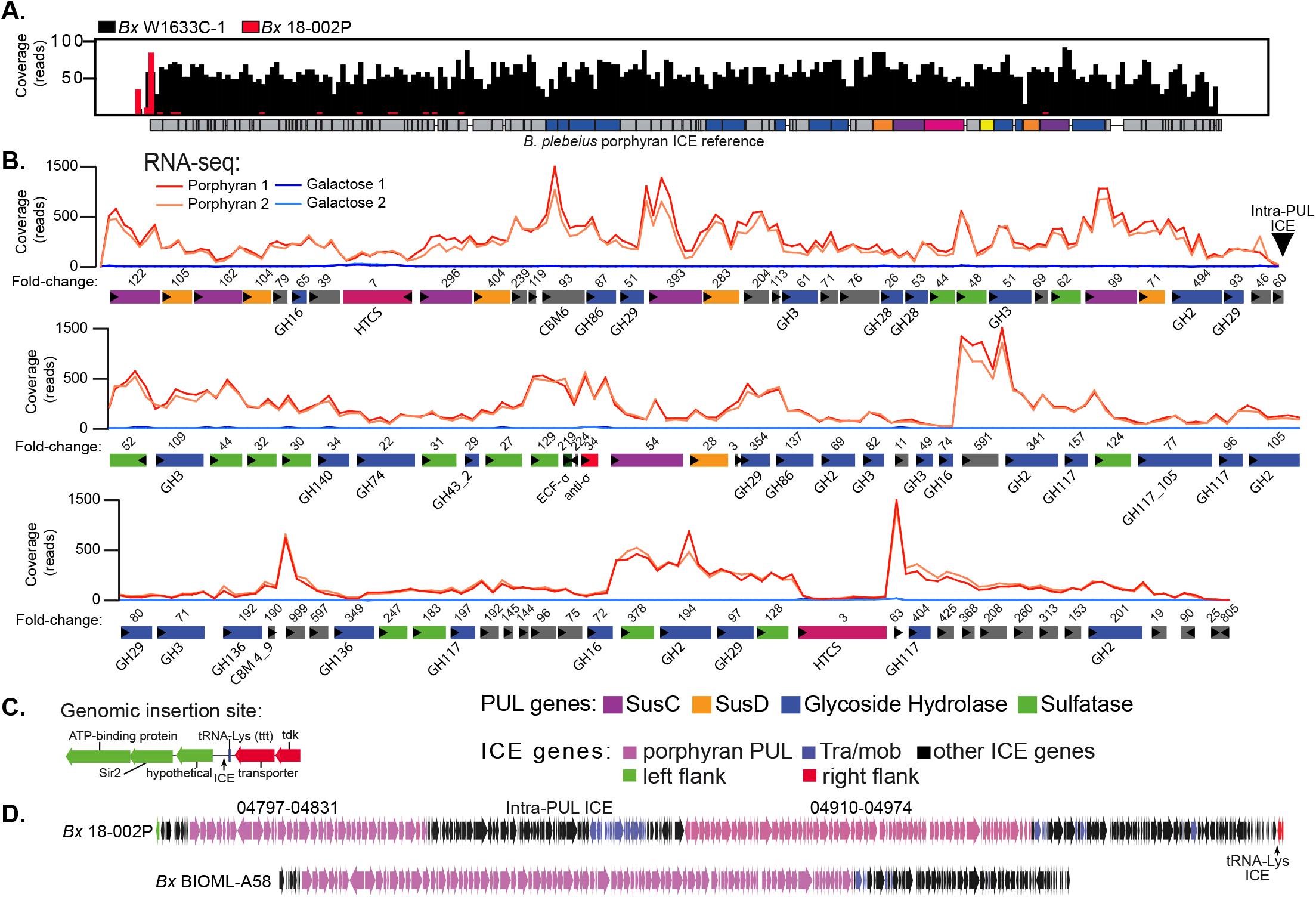
**(A)** Reference-guided mapping histograms to the *B. plebeius* porphyran ICE for two different strains: *Bx*^W1633C-1^ (black) and *Bx*^18-002P^ (red). *Bx*^W1633C-1^ has full coverage across the entire *Bp* ICE, while sequences for *Bx*^18-002P^ do not map to the known *B. plebeius* genes. **(B)** Transcriptional responses of genes within a 168.6kbp PUL in *Bx*^18-002P^ during growth on porphyran compared to galactose. **(C)** Higher resolution illustration of the genomic insertion site of the *Bx*^18-002P^ PUL. Comparative genomics against publicly available non-porphyran-degrading *Bx* genomes and subsequent PCR analysis (**Extended Data Figure 1**) demonstrated that the *Bx*^18-002P^ ICE is inserted downstream of a tRNA-lys^(TTT)^ and *tdk* gene. (**D**) Lower resolution comparison of the ICEs between the *Bx*^18-002P^ *and Bx*^BIOML-A58^ strains, highlighting the presence of extra genes, possibly the result of an additional ICE insertion into the middle of the *Bx*^18-002P^ PUL without disrupting its function.

The *Bx*^18-002P^ PUL also encodes additional GH families that have been shown to degrade other substrates suggesting that it encodes multiple glycan degrading capabilities. Using SACCHARIS^19^, putative enzymes within the *Bx*^18-002P^ locus were aligned with carbohydrate active enzymes with known functions. In addition to red algal galactans, this locus may target a diversity of substrates, which could include pectins (GH28s, GH140), ulvan (GH105), fucoidan (GH29), and β-mannans (GH2s). The presence of a β-galactofuranoidase (GH117), α-L-arabinofuranosidase (GH43), β-glucuronidase (GH2), and β-xylosidases (GH3s) suggests that this pathway may consume both red and green seaweed polysaccharides (**Supplemental Table 7**). Homo- and hetero-xylans are found in red and green seaweeds^20^, and complex xylogalacto(furano)arabinan is a primary component of edible green seaweed *Cladophora falklandica*^21^. Notably, however, there is a lack of polysaccharide lyases typically associated with the depolyermization of polyuronic acids found in these species. The collection of this diverse group of activities indicates that this pathway is tailored to hydrolyze structurally complex substrates found in edible seaweeds, and the presence of multiple regulatory proteins may assist in the fine-tuning of enzymatic cascades.

The carrageenan- (*Bt*^3731^, *Bo*^12C04^) and porphyran-degrading (*Bx*^18-002P^, *Bp*^17135^) strains encode several GH16 family enzymes characterized in the CAZy database as carrageenases, agarases and porphyranases, amongst others. Therefore, we used a GH16 family phylogeny as a proxy to explore GH family function into subfamilies, further demonstrating evolutionary trajectories towards substrate specificity. While the *Bt*^3731^ and *Bo*^12C04^ proteins remain tightly clustered in a separate clade, the *Bx*^18-002P^ and *Bp*^17135^ GH16 enzymes display broader subfamily assignments including subfamilies 12, 15 and 16 (**Extended Data 6**). This is further exemplified by the various oligosaccharides produced from different subfamilies of GH16 enzymes that allow for efficient depolymerization of seaweed glycans (**Extended Data 6**). These evolutionary adaptations towards glycan specificity undoubtedly play a critical role in proficient carbon utilization by these microbes but also provide an advantage for colonizing the human GI tract.

## Human gut Firmicutes with genetic upgrades to degrade seaweed polysaccharides

To date, all of the examples of seaweed-degrading human gut bacteria, including the isolates described above, have been Gram-negative Bacteroidetes. The Gram-positive Firmicutes that thrive in the human gut are typically more abundant, taxonomically more diverse and are also proficient fiber degraders. This raises the question of whether or not members of this phylum have received similar “genetic upgrades”. During studies with the same cohort of healthy human volunteers used to isolate seaweed-degrading *Bacteroides*, we fortuitously observed that some spore-forming gut bacteria (isolated by ethanol selection and re-germination on medium containing taurocholic acid^22^) had the property of pitting the agar plates on which they were cultivated. This phenotype is exhibited by the agarose and porphyran-degrading *Bacteroides* and suggested that these bacteria might possess the ability to degrade these substrates. From two separate people, we obtained isolates belonging to the same spore-forming species, *Faecalicatena contorta*, which have varying abilities to grow on agarose and/or porphyran (**Extended Data 3**). We generated high-quality, nearly closed genomes for each of these isolates (herein, *Fc*^548^ and *Fc*^588^) and performed RNA-seq during growth on porphyran or agarose relative to a galactose reference for each strain. Each strain’s transcriptional response during growth on seaweed polysaccharides was characterized by strong upregulation of a single gene cluster that was homologous between the two isolates (**Fig. 5A**, **Extended Data 7, Supplemental Tables 8, 9**). Based on homology to the genes identified, a third, related strain of *Faecalicatena fissicatena* (strain D5; herein, *Ff* ^D5^) that harbors the same locus was located in a public database. Growth and RNA-seq analysis of this strain on porphyran and agarose, confirmed the prediction that it possesses the ability to grow on these polysaccharides (**Extended Data 3,7; Supplemental Table 10**).

**Figure 5.**
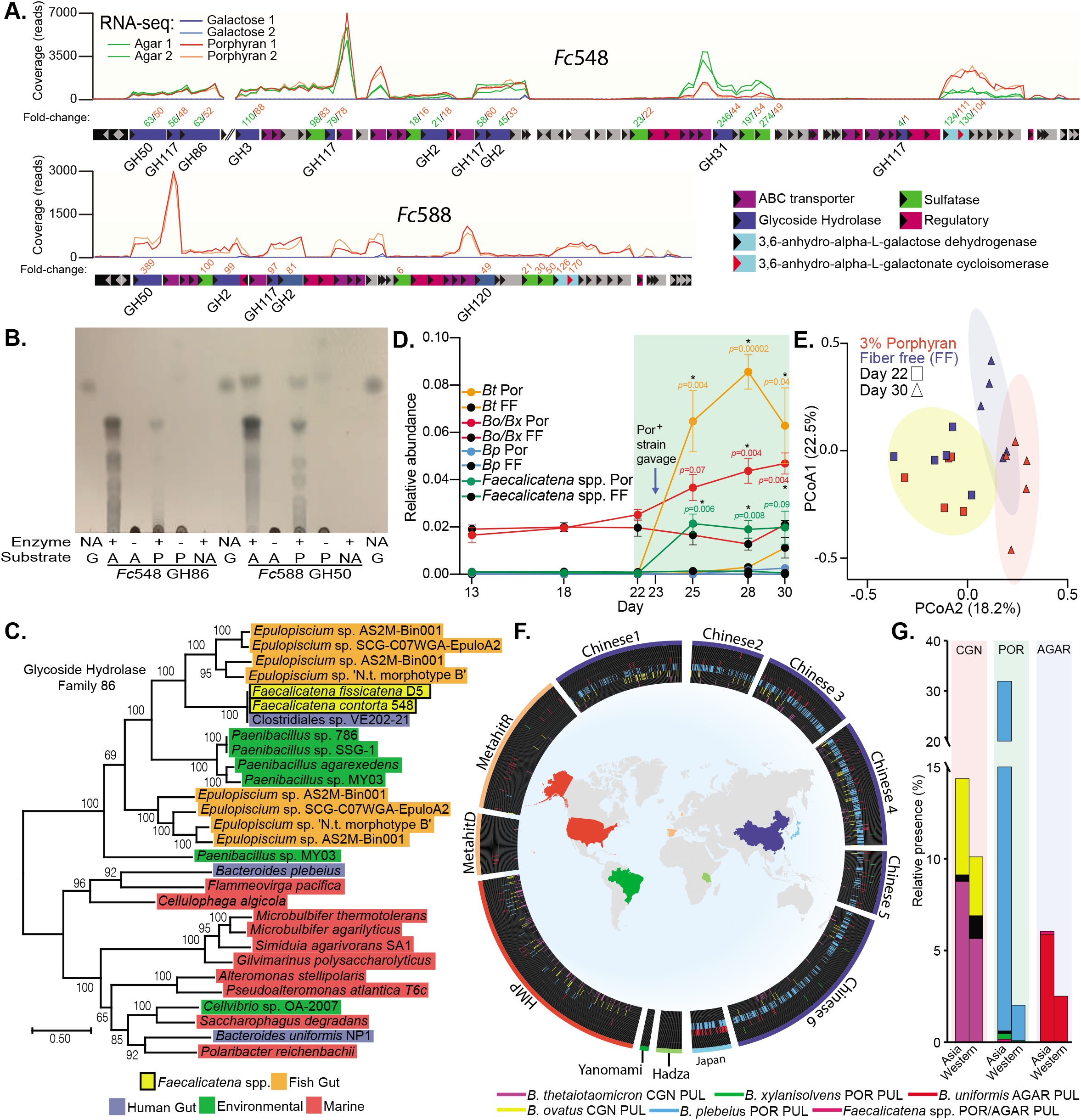
**(A)** Transcriptional responses of the homologous loci in *Fc*^548^ and *Fc*^588^ during growth on agarose and/or porphyran compared to galactose. **(B)** Thin-layer chromatography of *Fc*^548^ GH86 and *Fc*^588^ GH50 enzymes on high molecular weight agarose (A) and porphyran (P) with galactose (G) serving as a monosaccharide standard. **(C)** Phylogenetic position of *Faecalicatena* GH86 enzymes compared to other gut and environmental enzymes in this family, highlighting the similarity to enzymes found in *Epulopiscium* and *Paenibacillus* spp. **(D)** Engraftment of porphyran-degrading strains into gnotobiotic mice with a non porphyran-degrading human microbiota. Significant differences were observed in OTUs corresponding to *Bt*, *Bo/Bx* and *Faecalicatena* species when comparing the 3% porphyran diet (n=5) to a fiber free (FF) control diet lacking porphyran (n=5). Error bars are the standard error of the mean (S.E.M.) and unpaired *t*-tests were run using the Holm-Sidak correction to determine significance for days 25, 28 and 30 with significant *p*-values (<0.05, *) noted. **(E**) PCoA plot of individual mice shown in C before and after porphyran intervention in the presence of porphyran-degradung bacteria. (**F**) Global view of the distribution of known seaweed degrading gene clusters in existing metagenomic sequencing surveys. Colored wedges correspond to species respective PULs and outer band colors relate to country of data origin (**G**) Summary of CGN, porphyran and agarose degrading gene clusters in Western and Asian datasets based on the data shown in F.

The gene clusters identified in the three new *Faecalicatena* isolates encode predicted enzymes that belong to some of the same GH families as in agarose and porphyran-degrading *Bacteroides*. To determine if these enzymes are able to degrade these polysaccharides, we produced recombinant forms of 5 proteins from 3 different families (GH50, GH86, GH117). We observed activity with the GH50 (*Fc*^588^) and GH86 (*Fc*^548^) enzymes, which were capable of producing oligosaccharides from agarose and porphyran therefore suggestive of *endo*-acting function (**Fig. 5B**). However, three different GH117 enzymes—homologs of which are known to remove 3,6-anhydro-L-galactose from neoagarooligosaccharides^23,24^—displayed little detectable activity against either native substrates or GH86/GH50-derived oligosaccharides, indicating that their activity may require additional processing steps or recognition determinants to work (**Extended Data 8**).

The seaweed-degrading enzymes found in human gut *Bacteroides* have close relatives in marine Bacteroidetes, which are plentiful in the ocean and critical for turnover of algal biomass^25,26^. Interestingly, the enzymes contained in human gut *Faecalicatena* spp. share close relatives with marine Gram-positive bacteria, including *Paenibacillus* and *Epulopiscium* spp. (**Fig. 5C**, **Extended Data 9, Supplemental Tables 11, 12, 13**), the latter being abundant colonizers of marine surgeonfish^27^. Sequence-based surveys of surgeonfish gut microbiomes have previously detected enzymes in these same families in metagenome- and single cell isolation-based genome sequences belonging to *Epulopiscium* spp. Thus, it appears that the selective pressure imposed by human consumption of seaweed has been sufficient to not only select for multiple, independent transfers of agarose, carrageenan and porphyran utilization functions into gut *Bacteroides*, but also into human gut Firmicutes from a different genetic reservoir. Interestingly, each of the *Faecalicatena* agarose/porphyran loci was flanked by an integrase, which may have been involved in its mobilization (**Extended Data 7**). However, a lack of sequenced isolates in these two species that do not possess the ability to grow on agarose or porphyran prohibited us from reconstructing an ancestral genome state to test this hypothesis.

## Porphyran-mediated engraftment of a seaweed-degrading community in humanized mice

The presence of human gut *Bacteroides* with the ability to utilize polysaccharides like porphyran has catalyzed interest in using these relatively rare fiber-degrading abilities to engineer “orthogonal niches” in the gut microbiome^4,28^, which may be useful in implanting exogenous gut bacteria with engineered metabolism^4^. To determine if our expanded collection of human gut *Bacteroides* and *Firmicutes* could successfully colonize the gut in a dietary porphyran-dependent fashion, we colonized germfree mice (n=10 total) with feces from a single healthy human donor from which we repeatedly failed to enrich for indigenous agarose or porphyran-degrading bacteria. After 22 days of colonization with the human microbiota, we switched one group of 5 mice to a low fiber diet that was supplemented with 3% *Porphyra yezoensis* seaweed from food-grade unroasted nori sheets, while the other group was maintained on a nearly identical diet that lacked porphyran. On day 23 we gavaged both groups of mice with a community containing 8 different isolates (*B. plebeius*^17135^, *B. thetaiotaomicron*^PUR2.2^, *B. xylanisolvens*^18-002P^, *B. xylanisolvens*^W1633C-1^, *B. ovatus*^PUR1.1^, *F. contorta*^548^, *F. contorta*^588^, *F. fissicatena*^D5^) and maintained each group on their respective diets for another week (Day 30). Compared to the control group that was not fed *P. yezoensis*, those that were fed this culinary seaweed showed significant increases in the relative abundances of operational taxonomic units (OTUs) corresponding to *B. thetaiotaomicron* (4.6-fold higher) and *Faecalicatena* spp. (36.1-fold higher), which were two of the porphyran-degrading species that we introduced and did not have corresponding OTUs in the donor microbiome (**Fig. 5D**). In addition, *P. yezoensis* fed mice also showed an increase in an OTU corresponding to *B. ovatus/B xylanisolvens* (1.2-fold higher), which were also among the porphyran-degraders added. OTUs belonging to this group were already present in the transplanted human microbiome and remained ~2% before the intervention. After switching to the low fiber diet, or the matching diet containing *P. yezoensis, B. ovatus/B. xylanisolvens* decreased in abundance in mice fed the reference diet, which is consistent with the response of this fiber-degrading group in humanized mice fed low fiber diets^29^. However, in mice consuming *P. yezoensis*, *B. ovatus/B. xylanisolvens* showed the opposite trend and steadily increased over the one-week time period. Overall, feeding the *P. yezoensis* enriched diet, in concert with implanting seaweed-degrading bacteria, resulted in a significantly altered microbiome (**Fig. 5E**).

## Geographic distribution of seaweed degrading genes in human gut microbiomes

The close coupling of the gut microbiome to diet has made it possible for indigenous human gut bacteria to acquire new genes that enable them to compete for novel nutrients like seaweed polysaccharides. The extent to which these types of relationships have permeated the global microbiome remain to be revealed and may be impossible to determine without a more sophisticated appreciation of how many biochemically unique nutrients have been introduced into the human diet, often in geographically or culturally restricted fashions. However, we show here that genes for seaweed polysaccharide degradation have clearly penetrated the human gut multiple separate times and from different microbial gene reservoirs.

Since the selection for these genes is likely driven by regional dietary habits, we measured the presence of the known genes for seaweed degradation in global metagenomic surveys of human fecal samples. Consistent with several types of seaweed, including those that contain CGN, being consumed in parts of Asia for several millennia, the prevalence of these genes was substantially higher in metagenomic surveys of Chinese and Japanese subjects (**Fig. 5F,G**). The *B. plebeius-* type porphyran PUL was the most abundant, while the *B. uniformis*-type agarose PUL appeared to be enriched in the limited Japanese samples, often co-occurring with the *B. plebeius*-type PUL. The presence of the CGN ICE extended into the metagenomic samples from the United States, consistent with its use as a food additive and historical consumption in European populations, although prevalence appeared lower in European samples. While sampling has not yet been extended as deeply to populations living more traditional hunter-gatherer lifestyles, it is notable that none of these genes were detected in such populations from South America or Africa. Finally, the prevalence of the newly identified porphyran PUL and gene clusters from Gram-positive *Faecalicatena* spp. were both quite sparse in all samples. While detection of these genes will be influenced by sequencing depth, this did not hinder detection of the *B. plebeius*-type PUL, suggesting that species containing the latter genes are either more successful or more abundant. As metagenomic sequencing methods become more sophisticated and applied more broadly to global populations, parallel exploration of the novel genes that have permeated the human gut microbiome as a function of dietary selective pressure will reveal strategies for developing future engineered probiotics that can be coupled to short- or long-term term diet supplementation. Exploitation of these “orthogonal niches” holds powerful potential for engineering functions into the microbiome and implanting new microorganisms^4,28^. However, as evidenced by the unequal competition of the agarose/porphyran-degrading species that were implanted into humanized gnotobiotic mice, some of which had very similar genes, there is likely to be much left to understand about how these different gene clusters will function in different bacterial backgrounds and microbiomes.

## Methods

### Culture collection bacterial strains and growth conditions

A total of 354 human and animal gut Bacteroidetes were included in our initial survey. A complete list is provided in **Supplemental Table 1**, along with species designation based on 16S rDNA and associated meta-data. Species classifications were made based on BLAST alignment of each strain’s near full length 16S rRNA gene sequence to a database containing the type strains of the >40 named human gut Bacteroidetes species. Isolates with >98% 16 rDNA gene sequence identity to the type strain of a named species were labeled with that species designation. This classification strategy included all except 3 of the strains examined, which were all related to *B. uniformis* and had slightly more divergent 16S sequences (~96% identity to the type strain). Because of the small number of strains that did not satisfy our 98% cutoff, we grouped these unclassified strains with their nearest relative and labeled them as more divergent in **Supplemental Table 1**. All Bacteroidetes strains were routinely grown in an anaerobic chamber (Coy Lab Products, Grass Lake, MI) at 37°C under an atmosphere of 10% H_2_, 5% CO_2_, and 85% N_2_ on brain-heart infusion (BHI, Beckton Dickinson) agar that included 10% defibrinated horse blood (Colorado Serum Co.). *Faecalicatena* strains were grown in the same atmosphere on solid YCFA medium containing 0.1% taurocholic acid. For Bacteroidetes growth analysis, a single colony was picked into custom Chopped Meat Broth (CMB) and then sub-cultured into minimal medium (MM); For *Faecalicatena* strain growth analysis, CMB was used followed by a depleted medium (DM) (all media recipes can be found in **Supplemental Table 14**).

### Human subjects and enrichment for new seaweed degrading strains

Study participants were recruited through the Authentic Research Sections of the introductory biology laboratory course at the University of Michigan (BIO173). All participants gave written, informed consent prior to participating in the study. Participants under the age of 18 were granted permission by a parent or legal guardian. Participants ranged in age from 17 to 29, with a median age of 19. Individuals with self-reported history of inflammatory bowel syndrome, inflammatory bowel disease, or colorectal cancer were excluded from the study. This study was approved by the Institutional Review Board of the University of Michigan Medical School (HUM00094242 and HUM00118951) and was conducted in compliance with the Helsinki Declaration.

Human fecal samples were collected from healthy donors and grown in custom Chopped Meat Broth (CMB) in an anaerobic chamber (Coy Labs, Grass Lake, MI) with an 85% N_2_, 10% H_2_, 5% CO_2_ atmosphere. To isolate seaweed-degrading species, we transferred the CMB cultures (1:50) into minimal medium containing only one seaweed polysaccharide and incubated for ≥24 hrs. If growth was detected (defined as >0.1 OD_600_ above baseline, which was the same culture grown in minimal medium without added carbohydrates), cultures were streaked onto BHI plates containing 10% defibrinated horse blood (Quad Five, Ryegate, MT) and incubated for ≥48 hrs. Isolated colonies were sequentially purified by streaking 2-3x onto BHI-blood plates, then picked into fresh CMB and incubated for 24 hrs. Cultures were then grown in minimal media (MM) containing monosaccharides (MM-M) and incubated 24 hours prior to pelleting bacteria, washing twice in MM-no carbon (MM-NC) and transferring (1:100) into MM containing phenotypic respective seaweed polysaccharide. If isolates were still positive (*i.e*., based on the same >0.1 OD_600_ cutoff used above), isolates were stocked at −80°C and pelleted for DNA extractions. Kinetic growth assays were performed using 100ul 2X concentrated MM-NC mixed with 100 μL, 1 % carbohydrates (200 μL total culture volume containing 1:100 diluted and washed bacteria in 0.5% final carbohydrate) in 96-well plates (Costar) using a plate stacker coupled to a spectrophotometer (Biotek, Winooski, VT) as previously described^30^. The above isolation methods resulted in all *Bacteroides* species isolated in this study except the previously isolated *Bt*^3731^ and *Bo*^12C04^, which were part of existing collections and the former was isolated in the 1960s-1970s. However, *Faecalicatena contorta* 548 and 588 strains were fortuitously discovered by noticing a characteristic agarolytic “pitting” around colonies on YCFA agar plates. Elevated background growth in liquid YCFA-no carbohydrate led us to formulate a depleted medium (DM, described above) used for all *Faecalicatena* species analysis.

Low-melt agarose (Lonza, 50101) and carrageenan (Sigma, 22049, C1138 and 22048) were used for all experiments. Porphyran was prepared by the following method: Unroasted nori was purchased locally, ground in a blender and autoclaved for 3hrs at 50g/L in water. The solution was cooled to room temperature with stirring and crude filtered to remove debris. Porphyran was precipitated with 80% final volume ethanol overnight at 4°C. The precipitate was pelleted, supernatant removed and dissolved in water. The solution was centrifuged again to remove excess debris and co-extracted/pelleted polysaccharides (xylans). The porphyran containing supernatant was retained, lyophilized to reduce volume and extensively dialyzed against a 12-14 KDa dialysis membrane. Finally, the porphyran was lyophilized ad prepared at a 1% (10 mg/mL) autoclaved solution in water.

### DNA sequencing

*Bacteroides* DNA was isolated directly using the DNeasy Blood and Tissue Kit (Qiagen). For *Faecalicatena* isolates, a standard phenol-chloroform (1:1) method with bead-beating was employed. The 16S rDNA sequencing was performed at the University of Michigan DNA Sequencing Core with universal primers 8F (5’-AGAGTTTGATCCTGGCTCAG-3’) and 1492R (5’-GGTTACCTTGTTACGACTT-3’). Shallow draft genome sequencing of *Bacteroides* isolates was performed at the University of Michigan through the Host Microbiome Initiative Microbiome Core. However, strains with unknown genetic architectures, *Bx*^18-002P^, *Fc*^548^ and *Fc*^588^, were sequenced by MicrobesNG (Birmingham, UK) to obtain deeper, near-closed genome sequences. Reference based genome assemblies were generated using a 90% identity threshold in SeqMan NGen (DNASTAR) to determine coverage and generate corresponding read-mapping histograms. Short-read sequences (.fastq format) for each sequenced *Bacteroides* isolate were aligned against the known PUL or ICE sequences from *B. uniformis* NP1 (agarose), *B. plebeius* DSM17135 (porphyran), *B. thetaiotaomicron* VPI-3731 (carrageenan) and/or *B. ovatus* CL02T12C04 (carrageenan) loci involved in seaweed polysaccharide degradation. A corresponding reference guided variant analysis was also performed in SeqMan NGen to calculate percent nucleotide identity over covered regions to the respective ICE or PULS using 90% identity threshold, ≥50% depth variance and at least 3X sequencing coverage as cutoffs.

### RNA sequencing

Total RNA was extracted using a modified phenol-chloroform procedure to reduce seaweed polysaccharide carryover, namely for agarose and carrageenan, but not porphyran, which interfere with molecular biology applications by binding to RNA purification columns or forming solids. All strains were grown to near mid-log phase in 50 mL conical tubes and centrifuged to pellet bacteria. Cell pellets were washed with PBS to remove as much media as possible prior to resuspension in 3 mL RNA Protect Bacteria Reagent (Qiagen). After incubating for 5 min at room temperature (RT), cells were centrifuged at 7000xg for 10 min at RT, supernatants decanted and stored at −80°C until RNA extractions. Cell pellets were thawed on ice, resuspended in 200 μL TE (10mM Tris, 1mM EDTA, pH8.0) buffer with lysozyme (1mg/mL) and incubated at RT for 2 min prior to adding 700 μL RLT Buffer (Qiagen) with 10ul/mL B-mercaptoethanol (Sigma). After mixing thoroughly, 500ul phenol:chloroform:isoamyl alcohol (125:24:1; pH 4.5) was added and mixed by inversion. Following an 18,000xg centrifugation for 3 min at 4°C, the aqueous phase was removed into a new RNase free tube and another 500 μL phenol:chloroform:IAA was added, mixed and centrifuged as above. The aqueous phase was removed, 0.1 volume of 3M sodium acetate (pH5.5) and 600 μL cold 100% isopropanol were added and mixed by inversion. Importantly, tubes were rapidly centrifuged at 18,000xg for 1 min at RT to pellet precipitated polysaccharides. Some RNA may be lost at this stage but is required to remove residual seaweed polysaccharides (agarose and carrageenan) that interferes with downstream applications. Larger initial bacterial culture volumes (*i.e*., 25-100 mL) were required to overcome this loss. The RNA containing supernatant was retained and RNA was precipitated at −80°C for 20 min before centrifuging at 18,000xg for 20 min at 4°C. Supernatants were discarded and RNA pellets were washed with 70% isopropanol prior to centrifuging at 18,000xg for 5 min at 4°C. Supernatants were discarded and pellets were dried at RT before resuspending in 100 μL nuclease free water. RNA was subjected to DNaseI treatment (NEB) prior to re-precipitating and pelleting of RNA as above, including the polysaccharide removal centrifugation step. Finally, RNA pellets were washed with 1.3 mL 70% isopropanol, prior to drying and resuspension in 50 μL RNase free water. Total RNA was quantified using a Qubit 2.0 (Invitrogen).

To ensure a DNA-free RNA extraction was successful, quantitative polymerase chain reaction (qPCR) was performed on all RNA samples using a standard curve of genomic DNA and 16S rDNA primers for respective strains. All samples had <0.01ng/μL of 16S rDNA and proceeded to rRNA depletion using the MICROB*Express* Bacterial mRNA Enrichment Kit (Thermo Fisher) two times to sufficiently remove rRNA. Residual mRNA was quality controlled and converted to sequencing libraries at the University of Michigan Sequencing Core with an Illumina HiSeq-4000. Barcoded data were demultiplexed and analyzed using Arraystar software (DNASTAR, Inc.) using RPKM normalization with default parameters. Gene expression during growth on seaweed polysaccharides was determined by comparison to a galactose reference.

Genes with significant up-or down-regulation were determined by the following criteria: genes with an average fold-change >10-fold and both replicates with a normalized expression level >1% of the overall average RPKM expression level.

### Proteome analysis

For proteome analysis, *Bt*^3731^ was cultured in triplicate in MM with either 0.1% carrageenan from *Cladosiphon okamurans* (IEX purified) or 0.2% galactose, using previously described culture conditions^31^ Pre-cultures with the same carbon source were used to set the starting OD_600_ of the main culture to 0.05. Cells were harvested in mid-exponential phase (~0.25 OD_600_) by centrifugation at 4,000 x g for 10 min at 4°C and stored at −80°C until analysis. The soluble proteome (all soluble proteins), the secretome (all secreted proteins) and the membrane proteome (all proteins attached to membranes) were selectively enriched for further analysis. The soluble total proteome was extracted from cell pellets of carrageenan- and galactose-grown *Bt*^3731^ by sonication in TE buffer as previously described^32^. The secretome of carrageenan-grown cells was enriched from carrageenan culture supernatants using StrataClean beads^33^ and the membrane proteome was enriched from pellets of carrageenan-grown cells using the trypsin shaving approach^34^. Soluble proteome and secretome samples were further analyzed in a gel-based proteomic approach, i.e. protein extracts were separated on a 10% polyacrylamide sodium dodecyl sulfate mini gel as previously described^32^. After CBB staining, entire gel lanes were dissected into 10 equal pieces, which were destained and overnight-digested with trypsin (1 μg/mL, sequencing grade, Promega), before MS analysis of the peptide mixes. *Bt*^3731^’s secretome samples were subjected to gel-free digestion^33^ before MS analysis. Peptides were separated on a C18 column and subjected to reversed-phase chromatography on a nano-ACQUITY-UPLC (Waters Corporation). Mass spectrometry (MS) and tandem mass spectrometry (MS/MS) data were acquired using an online-coupled LTQ-Orbitrap mass spectrometer (Thermo Fisher)^34^. MS spectra were searched against a target-decoy protein sequence database, which included all predicted proteins of *Bt*^3731^ in forward and reverse directions (decoys) and a set of common laboratory contaminants. Validation of MS/MS-based peptide and protein identifications was performed with Scaffold v4 (Proteome Software Inc., Portland, OR, USA) using a maximum false discovery rate of 0.01 (1%) on the peptide level and 0.01 (1%) on the protein level. Only proteins that could be detected in at least two out of three biological replicates were considered identified. Normalized spectral abundance factors (%NSAF) were calculated samplefor each protein by normalizing Scaffold’s ‘total spectrum counts’ against protein size and against the sum of all proteins in the same. %NSAF values give a proteins percentage relative to total protein abundance in a given sample, thus allowing for comparisons between individual samples^35^.

### Carbohydrate-polyacrylamide gel electrophoresis (C-PAGE) of carrageenases

The genes encoding for carrageenases GH82a, GH82b and GH16 were amplified by PCR using gene-specific primers and cloned into a pET-28a(+) vector. In the resulting plasmids, GH82a was fused to an N-terminal His6 tag and GH82b, GH16 to C-terminal His6 tags. The signal peptide and few N-terminal sequences were removed to increase solubility. All the constructs were transformed into *E.coli* BL21 (DE3) cells. A single colony from each of these constructs was used to inoculate an LB culture containing 50 μg/mL kanamycin at 37°C overnight with shaking at 150 rpm, respectively. These starter cultures of GH82a and GH16 were used to inoculate 1L of auto-induction ZYP-5052 medium supplemented with 50 μg/mL kanamycin and incubated at 20°C for 4 days with shaking at 150 rpm. 1L LB containing 50 μg/mL kanamycin was inoculated with GH82b starter culture at 37°C until an OD_600_ of 0.8 was reached. IPTG at a final concentration of 0.5 mM was added to the LB culture to induce protein production at 16°C with overnight shaking at 150 rpm. Cells were harvested and stored at −80°C.

To purify enzymes, stored cells were first thawed and then lysed using chemical lysis. Tris(2-carboxyethyl)phosphine (TCEP) was added at a final concentration of 1 mM to prevent disulphide bridge formation during lysis. A soluble lysate containing expressed protein was obtained by centrifugation at 16,000 x g at 4°C for 45 minutes. The enzymes were purified by immobilized metal ion affinity chromatography (IMAC) using a Ni^2+^ loaded HiTrap HP column. Protein elution was achieved using 20 mM Tris-HCl pH 8 and 500mM NaCl buffer with an increasing imidazole gradient up to 500 mM concentration. Fractions were verified for successful protein expression using SDS-PAGE and dialysed overnight in 20 mM Tris-HCl pH 8, 1 mM DTT and 250 mM NaCl. Samples were concentrated (Amicon) using a 10 KDa cut-off membrane and stored at 4°C.

Carrageenase activity screening was performed by incubating enzymes GH82a, GH82b and GH16 with 3 different types of carrageenan: λ, κ and l-carrageenan at a final concentration of 0.3% at 37°C overnight. The enzymatic degradation products were analysed by carbohydrate-polyacrylamide gel electrophoresis (C-PAGE) and visualized using 0.005% Stains-All in 50% ethanol solution followed by destaining with 10% ethanol.

### Identification of closest homologs for genes involved in seaweed degradation

The amino acid sequence for each identified gene in CGN, agarose and porphyran degradation was subjected to a protein BLAST (NCBI) search against non-redundant Refseq to identify its closest relative and the corresponding environmental association of the bacteria in which that closest relative exists. We omitted results from the strains investigated in this study, and the bacterium with the highest match was investigated for its environmental source using NCBI BioProject information or PubMed literature search. Additionally, the organisms in which the closest homolog(s) resides were listed as Gram negative or positive as an additional indication of the genetic reservoir of the potentially transferred genes.

### Glycoside hydrolase 16 subfamily phylogeny

The GH16 enzymes from the *Bt*^3731^, *Bo*^12C04^, *Bx*^18-002P^, and *Bp*^17135^ were identified using dbCAN, and sequences were run through the SACCHARIS pipeline^19^. SACCHARIS combines user sequences with sequences from the CAZy database^36^ and trims sequences to the catalytic domain using dbCAN2^37^. Sequences were aligned with MUSCLE^38^, and a best-fit model was generated with ProtTest^39^. The final tree was generated with FastTree^40^ and visualized with iTOL^41^.

### *Faecalicatena* spp. amino acid phylogeny

FASTA amino acid files were run through the dBCAN2 server to annotate glycoside hydrolase 50, 86 and 117 families present in the *Faecalicatena* genomes. A multiple sequence alignment was generated for each glycoside hydrolase family using ClustalW in MUSCLE (EMBL-EBI) from amino acid sequences found in *Faecalicatena* spp. genomes, CAZy database, BLAST results or Protein Data Bank. The output file was converted to a MEGA file and a Maximum-Likelihood phylogeny created using a 100-resampling bootstrap analysis in MEGA X software.

### *Faecalicatena* spp. protein expression and activity

Putative agarases and porphyranases were analyzed for signal peptides, which were subsequently removed if present prior to PCR amplification. Purified PCR products were ligated into the pETite vector containing an N-terminal His6 tag (Lucigen) and transformed into HI-Control 10G cells. After a 1hr recovery in Luria-Bertani (LB) broth at 37°C, cells were plated onto LB plates supplemented with kanamycin (30 μg/mL) and incubated overnight at 37°C. The fidelity of the constructs was confirmed prior to transformation into TUNER *E. coli* cells and plated onto LB with kanamycin. A single colony was picked into LB broth with kanamycin and incubated at 37°C overnight. A 1L flask of Terrific Broth (TB) was inoculated with 5 mL of overnight culture and incubated at 37°C with shaking at 200rpm until mid-log growth (0.6 OD_600_) when it was placed on ice. After 1hr on ice cells were induced with 0.2 mM final isopropyl-d-1-thiogalactopyranoside (IPTG). Cells were incubated with 200rpm shaking at 16°C overnight then pelleted and stored at −80°C until purification.

Enzymes were purified by centrifuging thawed cells in Talon Buffer (20 mM Tris, 300 mM NaCl, pH8.0) at 8,000 x g post sonication and purified on a HisPur Cobalt Resin (Thermo Scientific). Two separate elutions (5 mL (E1) and 10 mL (E2)) were performed and run on an SDS-PAGE gel to verify expression/purification. Soluble, expressed proteins were dialyzed overnight using a 12-14KDa membrane in Talon buffer. Enzyme activity assays on agarose and porphyran were done in Talon Buffer with 1 μM of enzyme and incubated overnight at 37°C. Assays were heat inactivated and subjected to thin-layer chromatography (TLC) to check for enzymatic activity.

### Gnotobiotic mouse colonization

Ten 8-week old C57BL/6NCrl germ-free mice were gavaged with a healthy human microbiota that was repeatedly screened to be negative for the ability to enrich agarose or porphyran degrading bacteria. All mice were kept on normal chow for 13 days prior to switching to a fiber-free diet (TD.130343, Envigo). At day 22, 5 mice were switched to a diet containing 3% *Porphyra yezoensis* (TD.190608, Envigo), which was ground into a course powder and added to TD.190608 to replace 3% of the dextrose contained in the base diet. To investigate engraftment of the different species tested, equal volumes of *B.plebeius*^17135^, *B. thetaiotaomicron*^PUR2.2^, *B. xylanisolvens*^18-002P^, *B. xylanisolvens*^W1633C-1^, *B. ovatus*^PUR1.1^, *F. contorta*^548^, *F. contorta*^588^, *F. fissicatena*^D5^ were gavaged into all mice at day 23. After 8 days of 3% *Porphyra yezoensis* diet feeding in the experimental group (n=5) and 7 days post porphyran-degrading strain gavage (n=10), all mice were sacrificed. Feces were collected from each mouse throughout the experiment at the time points shown in Fig. 5.

For DNA extractions, fecal pellets were combined with acid-washed glass beads (212-300 μm; Sigma-Aldrich, USA), 500 μL Buffer A (200 mM NaCl, 200 mM Tris, 20 mM EDTA), 210 μL SDS (20% w/v, filter-sterilized) and 500 μL phenol:chloroform (1:1). A Mini-BeadBeater-16 (Biospec Products, USA) was used to disrupt the bacterial cells for 3 min at room temperature then centrifuged and the aqueous phase was recovered. An equal volume of phenol:chloroform (1:1) was added to the aqueous phase and was mixed with the aqueous phase by gentle inversion. After centrifugation (12,000 rpm, 4°C, 3 min), the aqueous phase was recovered and 500 μL of pure chloroform was added to the aqueous phase, mixed by inversion and the tubes were centrifuged (12,000 rpm, 4°C, 3 min). The aqueous phase was transferred into fresh tubes and 1 volume of −20°C chilled 100% ethanol and 1/10 volume 3 M sodium acetate (pH 5.2) were added to the aqueous phase. The samples were mixed by gentle inversion and incubated at −80°C for 30 min, centrifuged for 20 min (12,000 rpm, 4°C) and the supernatants were discarded. The pellets were washed in 70% ethanol, air-dried and then resuspended in nuclease-free water. The resulting DNA extracts were purified by using DNeasy Blood & Tissue Kit (Qiagen).

The V4 region of the 16S rDNA gene was sequenced at the University of Michigan Microbial Systems and Molecular Biology Laboratory using an Illumina MiSeq platform as previously described^42^. The resulting16S rRNA abundance data were processed using the Mothur software package to reduce sequencing errors and remove chimeras as previously described^43^. Sequences were aligned to the SILVA 16S rRNA sequence database^44^. Sequences were clustered into operational taxonomic units (OTUs) using a 99% similarity cutoff. The R package ‘vegan’ was used to calculate and plot the principal coordinates analysis (PCoA) from the Bray-Curtis dissimilarity index based on phylotype classification of the community. Analysis of molecular variance (AMOVA) was used to determine significance between community structure differences of different groups of samples. The generated OTUs were analyzed by BLAST to identify the corresponding species. The average relative abundance for highly represented OTUs were generated for each treatment group and sampling date and fold-change differences calculated only at day 30. Unpaired *t*-tests were run in Prism (GraphPad) using the Holm-Sidak correction to determine significance of OTU changes by day.

### Survey of human metagenomic data sets

Available cohorts of human gut metagenomic sequence data (National Center for Biotechnology Information projects: PRJNA422434^45^, PRJEB10878^46^, PRJEB12123^47^, PRJEB12124^48^, PRJEB15371^49^, PRJEB6997^50^, PRJDB3601^51^, PRJNA48479^52^, PRJEB4336^53^, PRJEB2054^54^, PRJNA392180^55^, and PRJNA527208^56^ were searched for the presence of CGN/POR/AGAR degrading PUL nucleotide sequences from *Bt*^37371^, *Bo*^12C04^, *Bp*^17135^, *Bu*^NP1^, *Bx*^18-002P^ and *Faecalicatena* spp. using the following workflow: Each PUL nucleotide sequence was used separately as a template and then magic-blast v1.5.0^57^ was used to recruit raw Illumina reads from the available metagenomic datasets with an identity cutoff of 97%. Next, the alignment files were used to generate a coverage map using bedtools v2.29.0^58^ to calculate the percentage coverage of each sample against each individual reference. We considered a metagenomic data sample to be positive for a particular PUL if it had at least 70% of the corresponding PUL nucleotide sequence covered.

### Data accessibility

Shallow draft genome data of all isolates can be accessed using BioProject PRJNA625I5I and corresponding BioSamples SAMN14593814-45. *Bx*^18-002P^, *Fc*^548^ and *Fc*^588^ WGS data can be accessed using accession numbers XXX, XXX and XXX, respectively. All RNA-sequencing data can be accessed using GEO accession project number GSE149357 and corresponding samples GSM4498559-82. Mass spectrometry proteomics data have been deposited to the ProteomeXchange Consortium via the PRIDE^59^ partner repository with the dataset identifier PXD019149. During the review process, the data set can be accessed at https://www.ebi.ac.uk/pride/login with username: and password: MHEl6KP7.

## Supporting information

Merged Supplemental Tables 1-14

## Acknowledgements

We would like to thank Laurie E. Comstock for providing *B. ovatus* CL02T12C04 and Emma Allen-Vercoe for providing *F. fissicatena* D5 and the University of Michigan Germfree Mouse facility. We thank Robert Glowacki, Ana Luis, Matt Ostrowski, Sadie Gugel, Jaime Fuentes, Shaleni Singh, Darryl Jones and the Nicole Koropatkin Lab for critical feedback during this project. A grant from the Howard Hughes Medical Institute (HHMI Grant # 52008119) and support from the University of Michigan’s Host-Microbiome Initiative to TMS supported work with the human cohort. This work was supported by NIH grants (GM099513 and DK096023 to E.C.M). We appreciate Jana Matulla’s and Sebastian Grund’s assistance during proteome sample preparation and MS measurements, respectively. The work of SM, TS, DB and JHH was financially supported by grants (BE 3869/4-2, SCHW 595/10-2, HE 7217/2-2) of the Deutsche Forschungsgemeinschaft (DFG) in the framework of the research unit FOR 2406 “Proteogenomics of Marine Polysaccharide Utilization” (POMPU). FU was supported by a scholarship from the Institute of Marine Biotechnology e.V. An Agriculture and Agri-Food Canada grant (J-002262) supported work from DWA.

## Author contributions

NAP and KU tested Bacteroidetes strains for growth on polysaccharides and processed the data. NAP, AC, RA, AA, DM and TMS performed enrichment analysis from human fecal samples and 16S rDNA identification. NAP, YX, AA performed follow up growth analysis on isolated strains. NAP, GVP, DNB generated and analyzed genomic sequence data. NAP, GP performed and analyzed RNAseq data. GVP performed the metagenomic analysis. NAP, MC and JP performed recombinant enzyme studies. NAP, GVP, JHH, ECM wrote the paper. NAP, DWA and JT performed phylogenetic analysis of enzymes. NAP and GVP performed the *in vivo* colonization experiment. All authors contributed to paper editing and agreed on the final draft. MC, FU, SM and TS conducted the proteome analysis for which DB provided resources.

## Extended Data Figure Legends

**Extended Data Figure 1.**
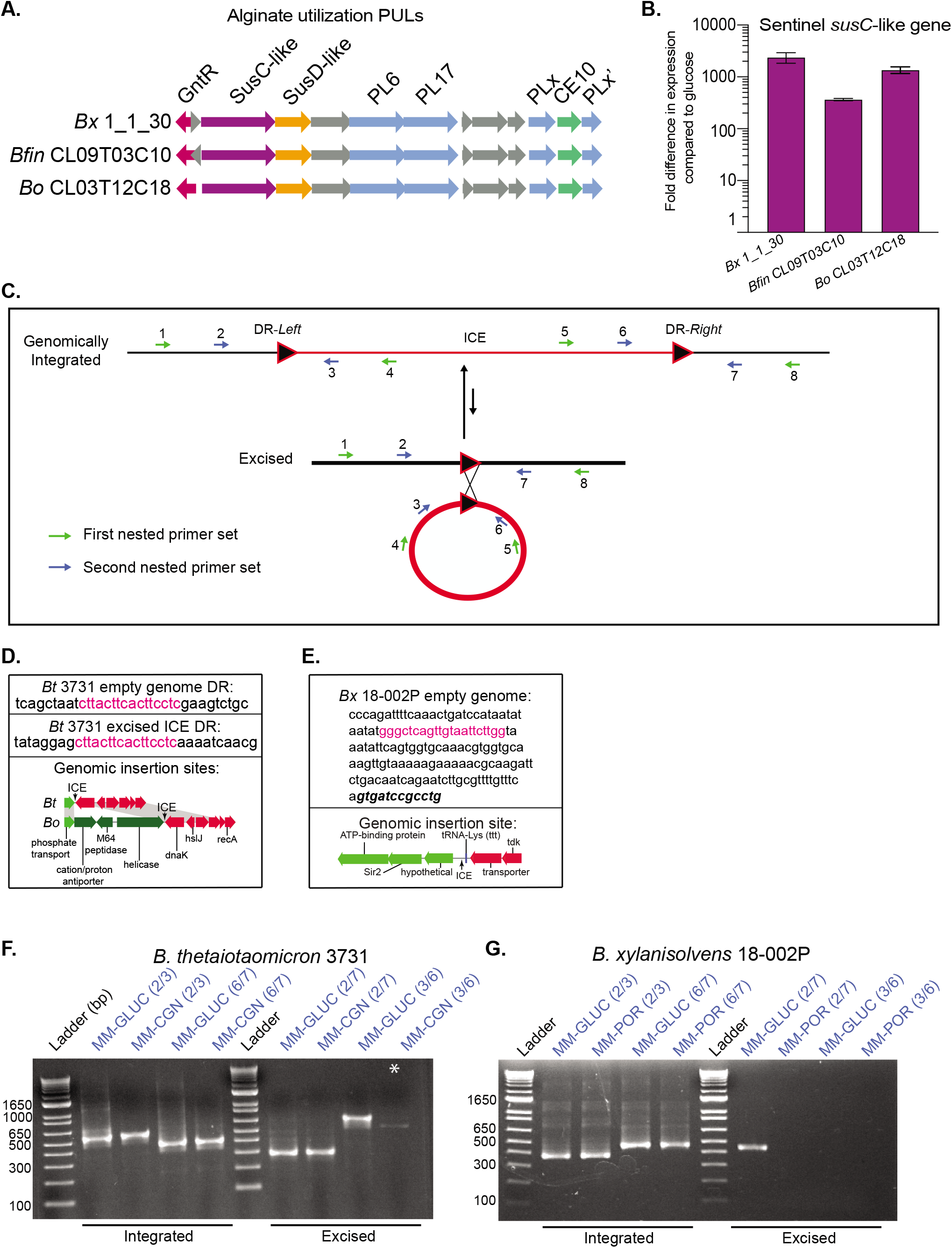
**(A)** Polysaccharide utilization loci (PULs) involved in alginate utilization from strains of the 3 different species we found to utilize alginate in this study. **(B)** Expression of alginate responsive sentinel *susC*-like genes for representative strains by qRT-PCR compared to glucose as a reference. **(C)** A schematic of the nested PCR approach used to examine possible ICE excision. **(D)** DNA sequencing results and corresponding genomic insertion site schematic for *Bt*^3731^ with pink nucleotides highlighting the direct repeat that presumably mediates insertion. **(E)** DNA sequencing results and corresponding genomic insertion site schematic for *Bx*^18-002P^ with pink nucleotides highlighting the direct repeat that presumably mediates insertion. **(F)** Agarose gel showing the PCR products from the second round of the nested PCR for *Bt*^3731^ where both empty genome and excised products are produced indicative of ICE excision. Note that the prominent band in the last well (MM-CGN (3/6 primer set)) is a PCR artifact and not the excised ICE form (marked with *). **(G)** Agarose gel showing the PCR products from the second round of the nested PCR for *Bx*^18-002P^ where an empty genome product is formed indicating some level of ICE excision.

**Extended Data Figure 2.**
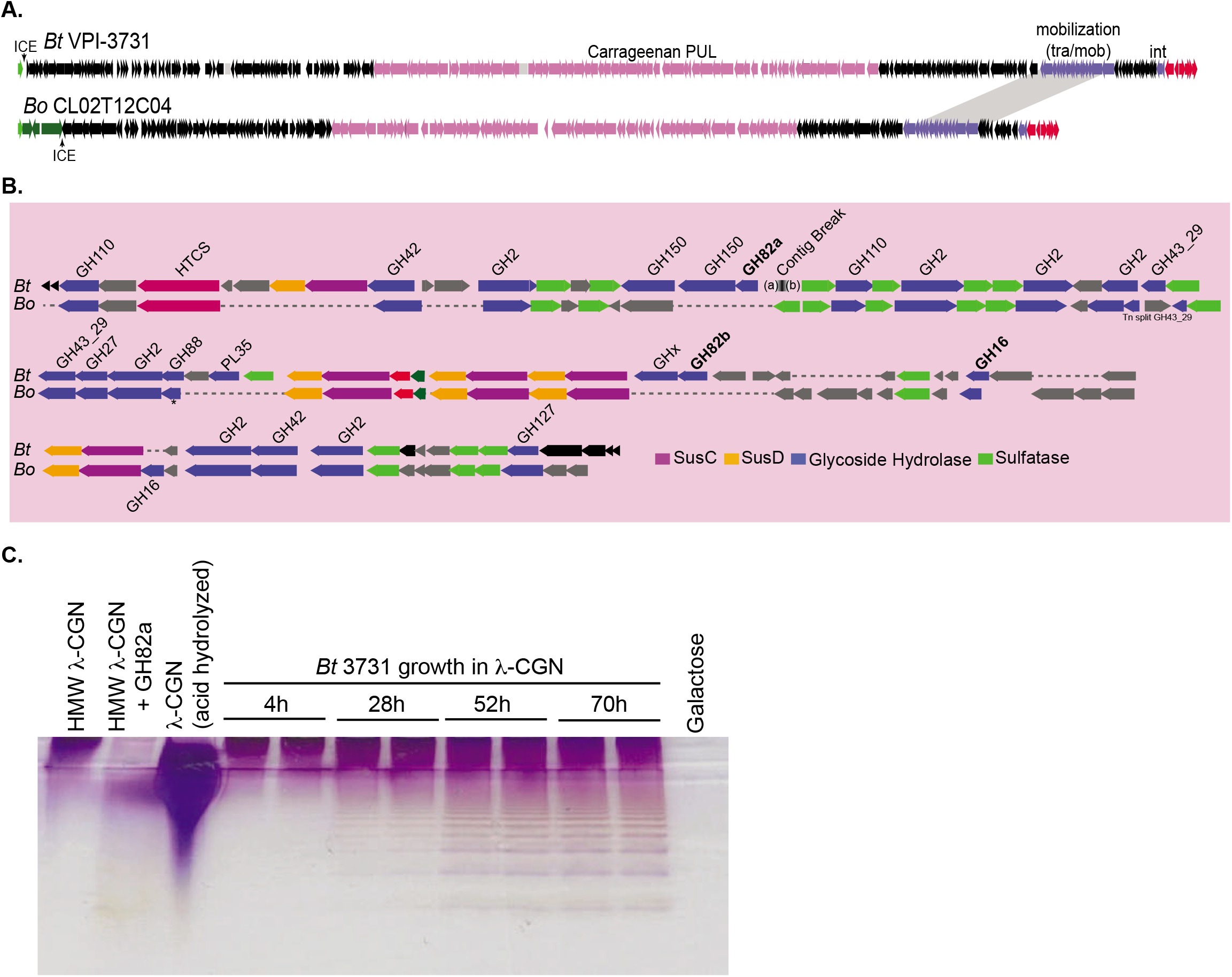
**(A)** Lower resolution schematics of the *Bt*^3731^ and *Bo*^12C04^ CGN-ICEs and flanking integration sites. **(B)** Higher resolution schematics of the *Bt*^3731^ and *Bo*^12C04^ CGN-PULs highlighting the genes that are syntenic and/or homologous between the two loci. The dashed lines are included to align PUL genes and do not correspond to actual sequence. **(C)** C-PAGE analysis of production of lower molecular weight “poligeenan” oligo- and polysaccharides produced over time during *Bt*^3731^ growth (n=2) on 0.2% λ-CGN. The poligeenan pattern observed is similar to the product profile of GH82a digested λ-CGN as compared to untreated and acid hydrolysed λ-CGN. The last lane shows supernatant of *Bt*^3731^ grown for 52h in galactose medium to control for capsular polysaccharide or other products made by this strain. Gel was stained overnight using 0.005% Stains-All in 50% ethanol solution followed by destaining in 10% ethanol.

**Extended Data Figure 3.**
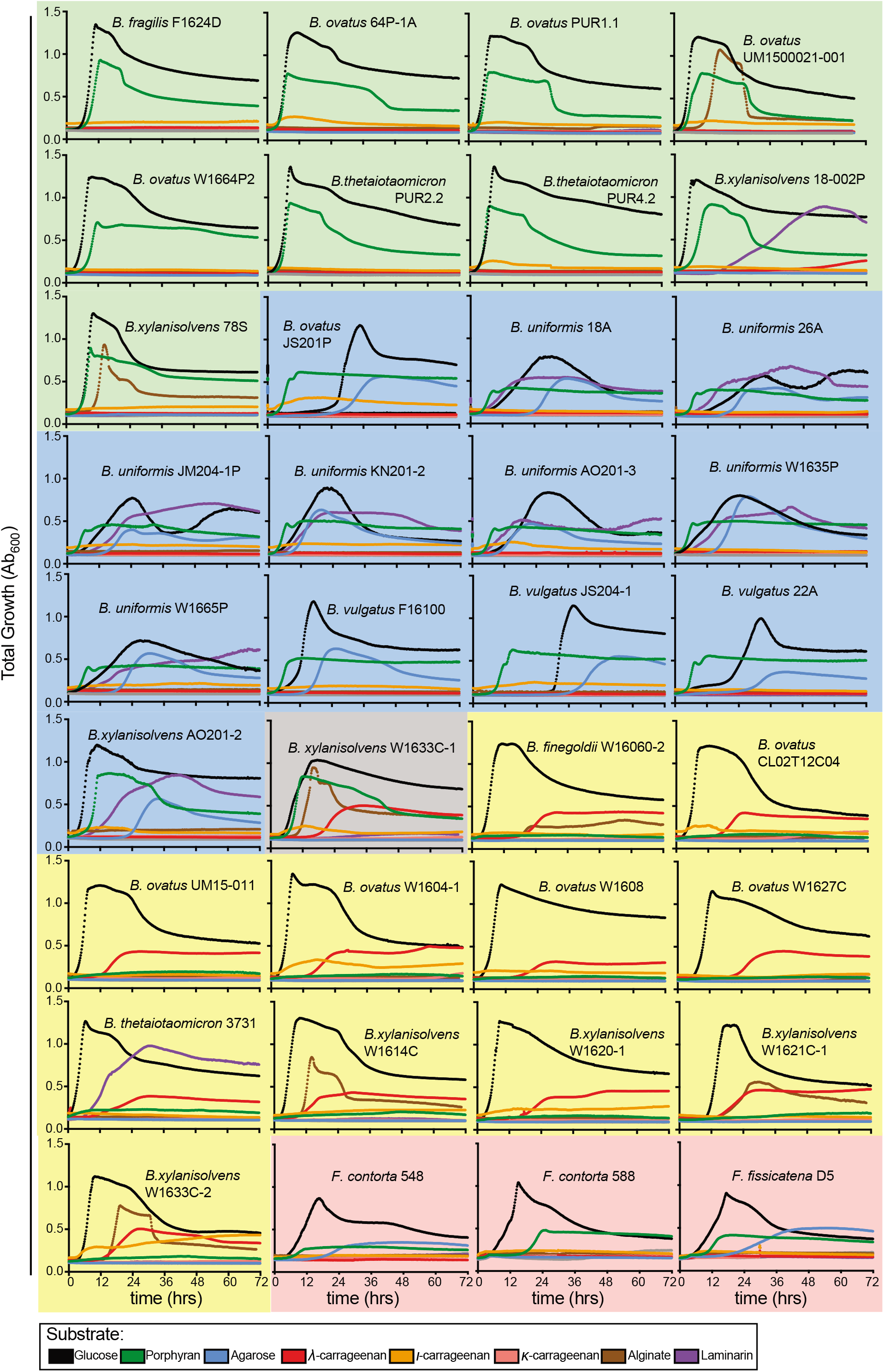
Average growth curve analyses (n=6) of seaweed degrading bacterial isolates on 7 different substrates, plus glucose as a control. The green background is for porphyran only degraders, blue for agarose-porphyran degraders, yellow for carrageenan and red for the Gram positive *Faecalicatena* species. The gray background is for *Bx*^W1633C-1^ that has both porphyran and carrageenan degrading capabilities.

**Extended Data Figure 4.**
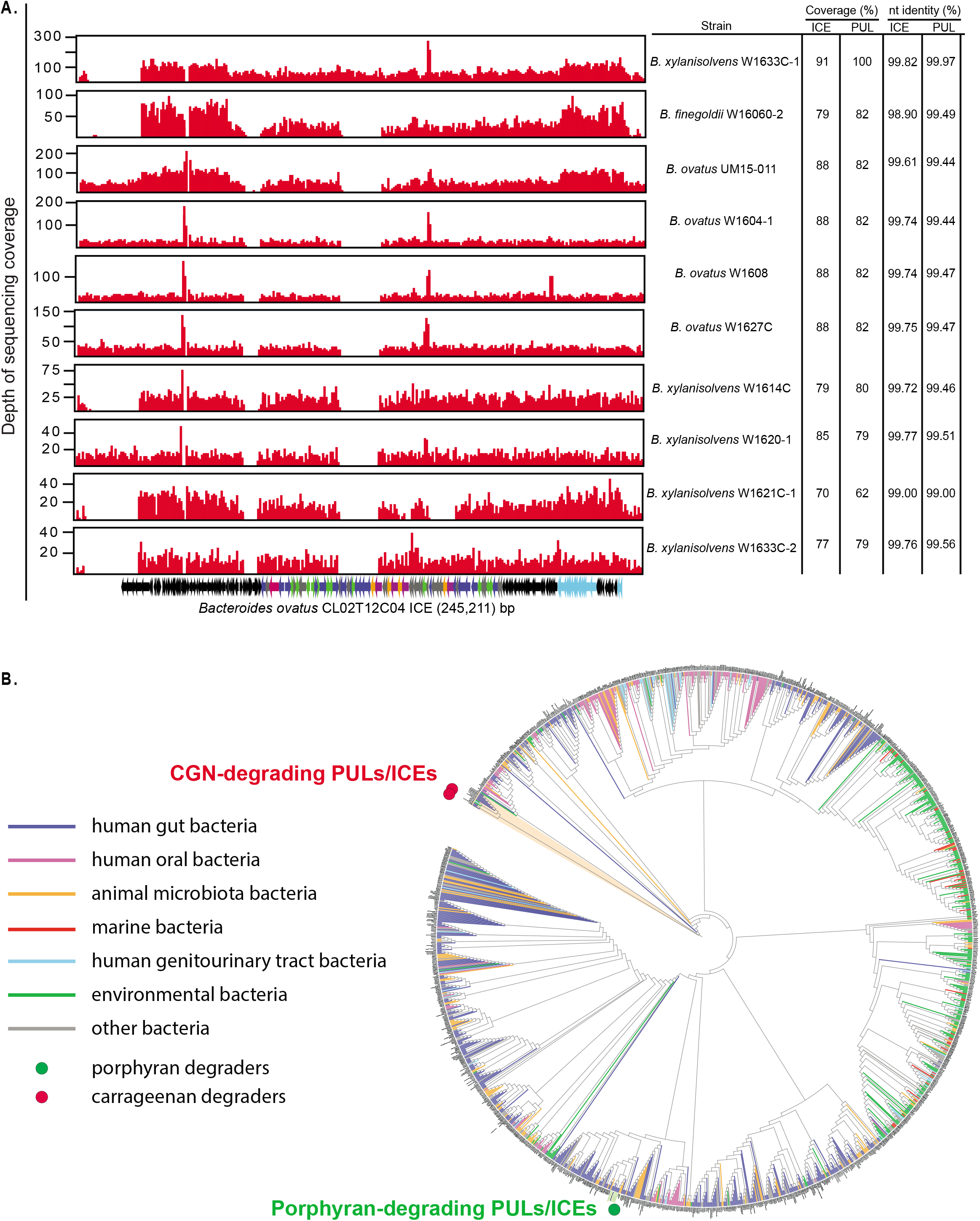
**(A)** Histograms showing reference guided mapping of CGN-degrading strain genomes to the *Bo*^12C04^ CGN-ICE template. The ICE and PUL linear coverage and nucleotide identity of covered regions of the ICE and PUL are shown as percent identity on the right. **(B)** A tree of 2,255 unique Bacteroidetes TraJ protein sequences from publicly available data and the genomes reported here. The locations of the TraJ sequences associated with the *B. plebeius*-type porphyran ICE and the newly identified CGN ICE are highlighted with green and red circles, respectively.

**Extended Data Figure 5.**
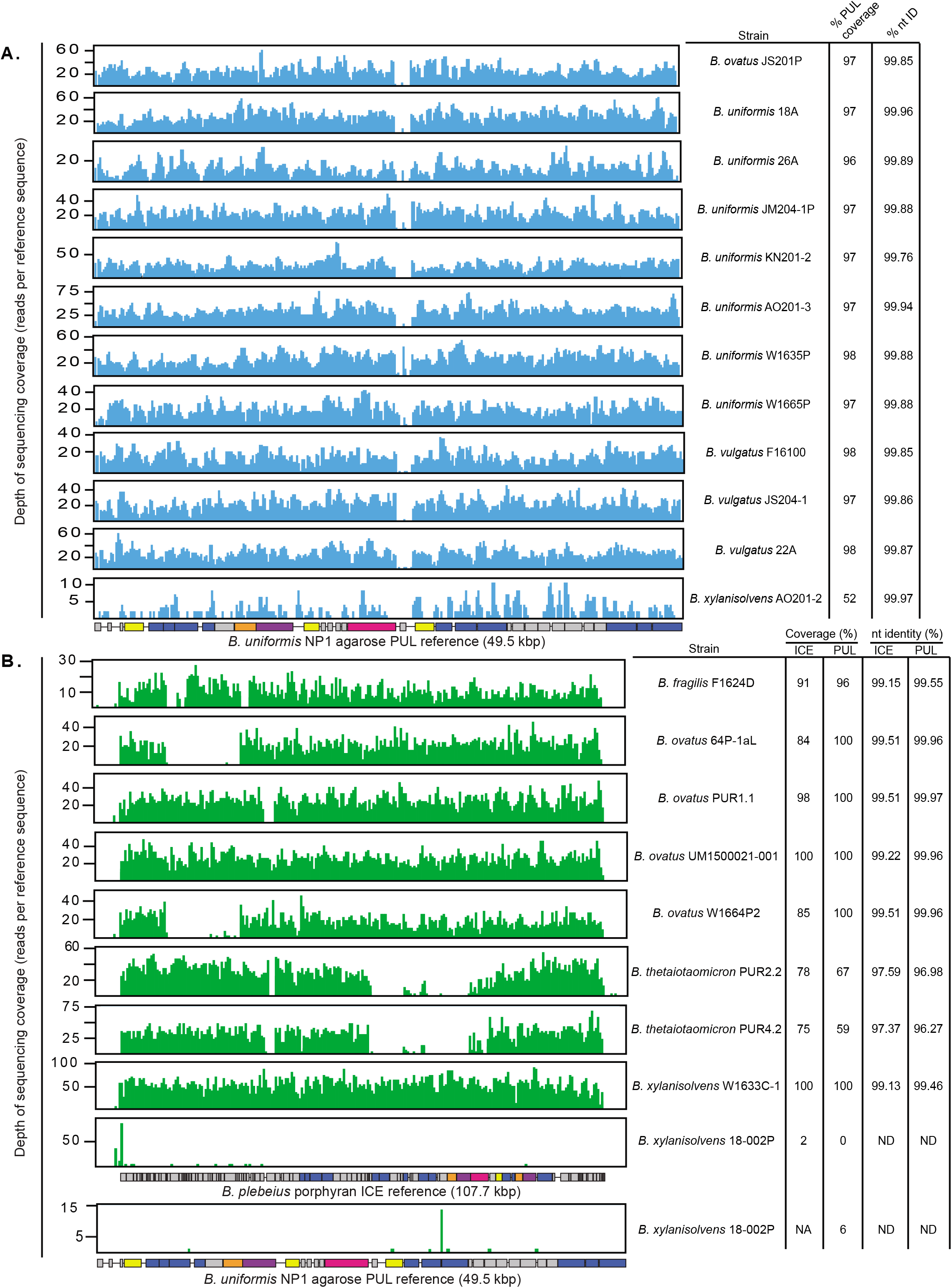
**(A)** Mapping of genomic reads from agarose-porphyran degrading strains to the *Bu*^NP1^ reference PUL showing linear coverage across the genomically inserted PUL and percent of coverage and nt identity. **(B)** Mapping of genomic reads from porphyran degrading strains to the *Bp*^17135^ reference PUL showing linear coverage across the ICE and percent of coverage and nt identity. Mapping of the *Bx*^18-002P^ assembly against the *Bu*^NP1^ reference is included at bottom to show the lack of homology to either of the known agarose or porphyran-degrading loci.

**Extended Data Figure 6.**
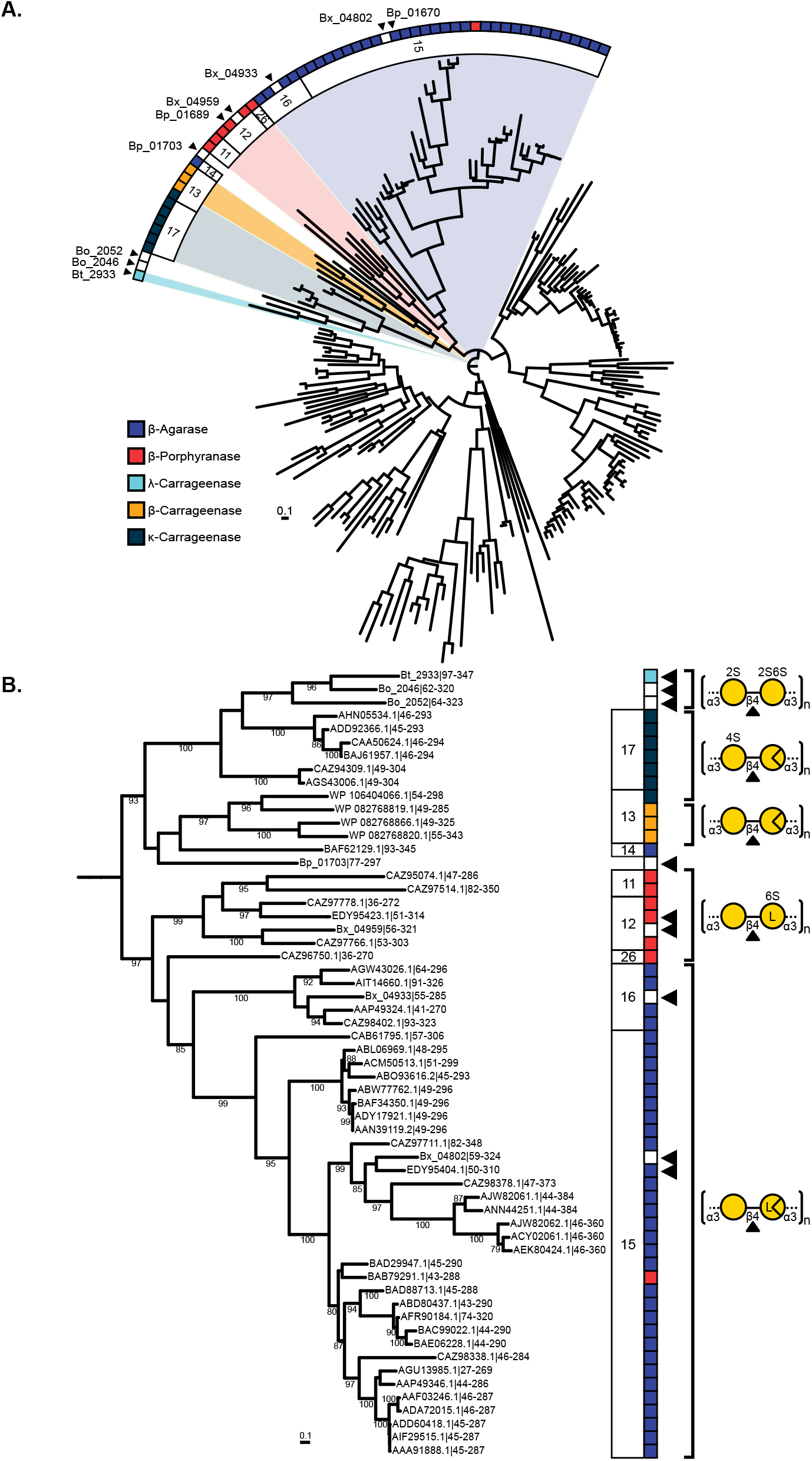
**(A)** Phylogenetic comparison of GH16 enzymes from the *Bt*^3731^ and *Bo*^12C04^ CGN PULs, and the *Bx*^18-002P^ and *Bp*^17135^ POR PULs against biochemically characterized GH16 members in the CAZy database. *Bt*, *Bo*, *Bx*, and *Bp* sequences are marked by black arrows. Characterized sequences of “red seaweed degrading” GH16 subfamilies (11-17, 26) are highlighted and members are colored by enzyme activity. **(B)** The above tree is pruned to display accession numbers and illustrate products from glycosidic cleavage.

**Extended Data Figure 7.**
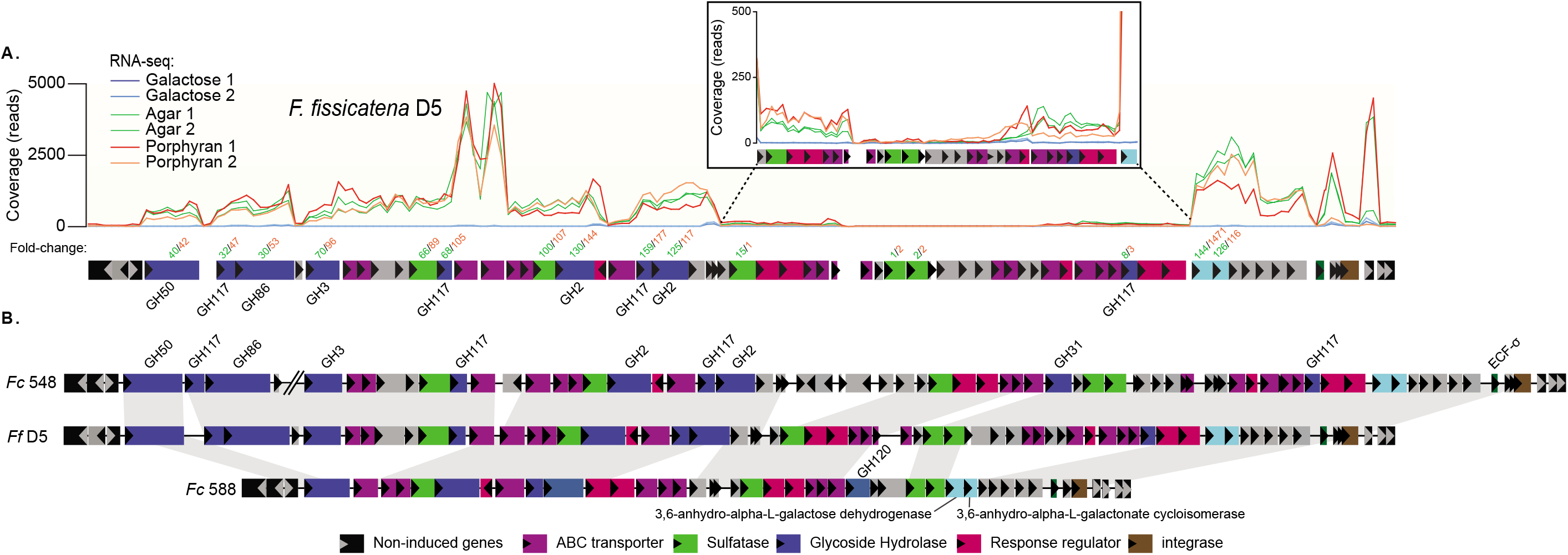
**(A)** Schematic of the gene clusters that are induced during growth on agarose and porphyran in *Faecalicatena fissicatena* D5 by RNA-seq when compared to galactose as a reference. The inset box shows a closer view of a section of genes that are induces less compared to their surrounding neighbors, but still show a positive response compared to galactose. **(B)** A genomic comparison of the three *Faecalicatena* spp. locus maps showing synteny and variation among the genes responsible for degrading agarose and/or porphyran. Note the first three enzymes in *Fc*^548^ are separated genomically from the rest of the genes in the loci by 110 genes (~87.9 kbp).

**Extended Data Figure 8.**
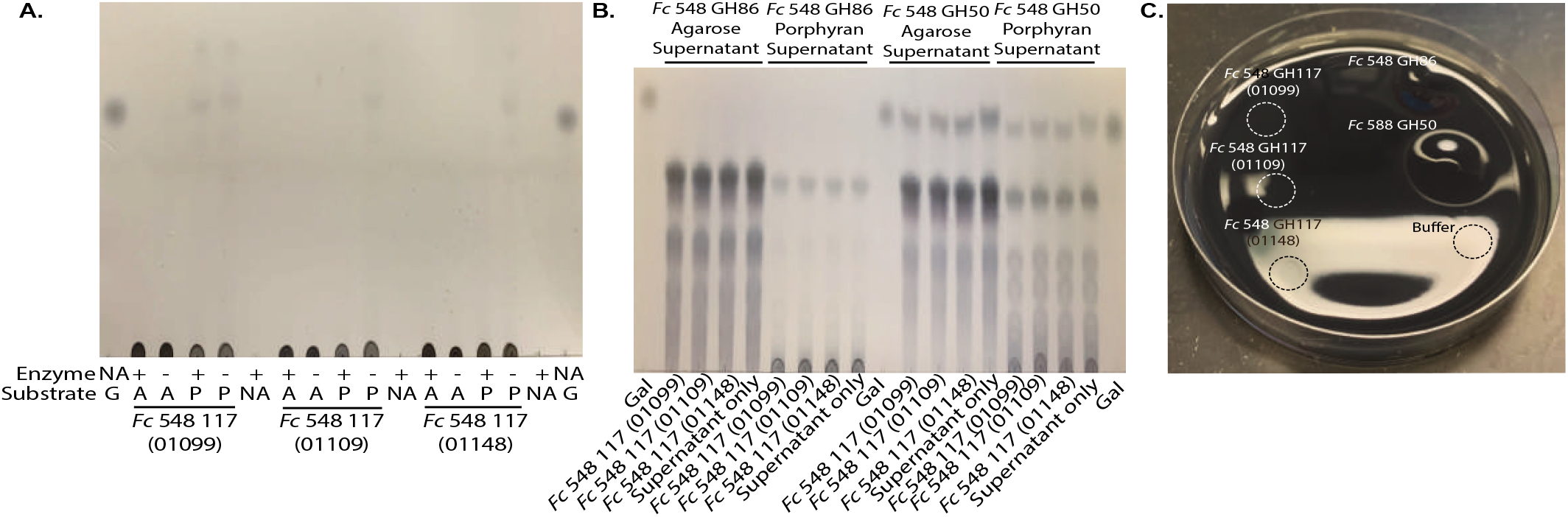
**(A)** Thin-layered chromatography analysis of enzymatic reactions for 3 of 4 GH117 enzymes found in *Fc*^548^. Minimal enzymatic activity is observed on agarose (A) and porphyran (P) substrates with galactose (G) as a size/sugar reference. These enzymes are often characterized as neoagarooligosaccharide hydrolases targeting smaller agarose fragments. The fourth GH117 found in *Fc*^548^ is 100% similar by amino acid sequence to one analyzed here and was omitted from analysis. **(B)** Thin-layered chromatography of enzymatic reactions of the same GH117 enzymes in A, but incubated with the supernatants of the GH86 (*Fc*^548^) and GH50 (*Fc*^588^) reactions. No further degradation of the agarose or porphyran oligosaccharides is observed after GH117 treatment. **(C)** An agar plate where the filter-sterilized recombinant glycoside hydrolases from *Faecalicatena contorta* species were spot plated and incubated overnight at 37°C. As observed with colonies growing on the same plates, the GH50 and GH86 display “pitting” even as pure enzymes, whereas the GH117 enzymes show little, if any, activity.

**Extended Data Figure 9.**
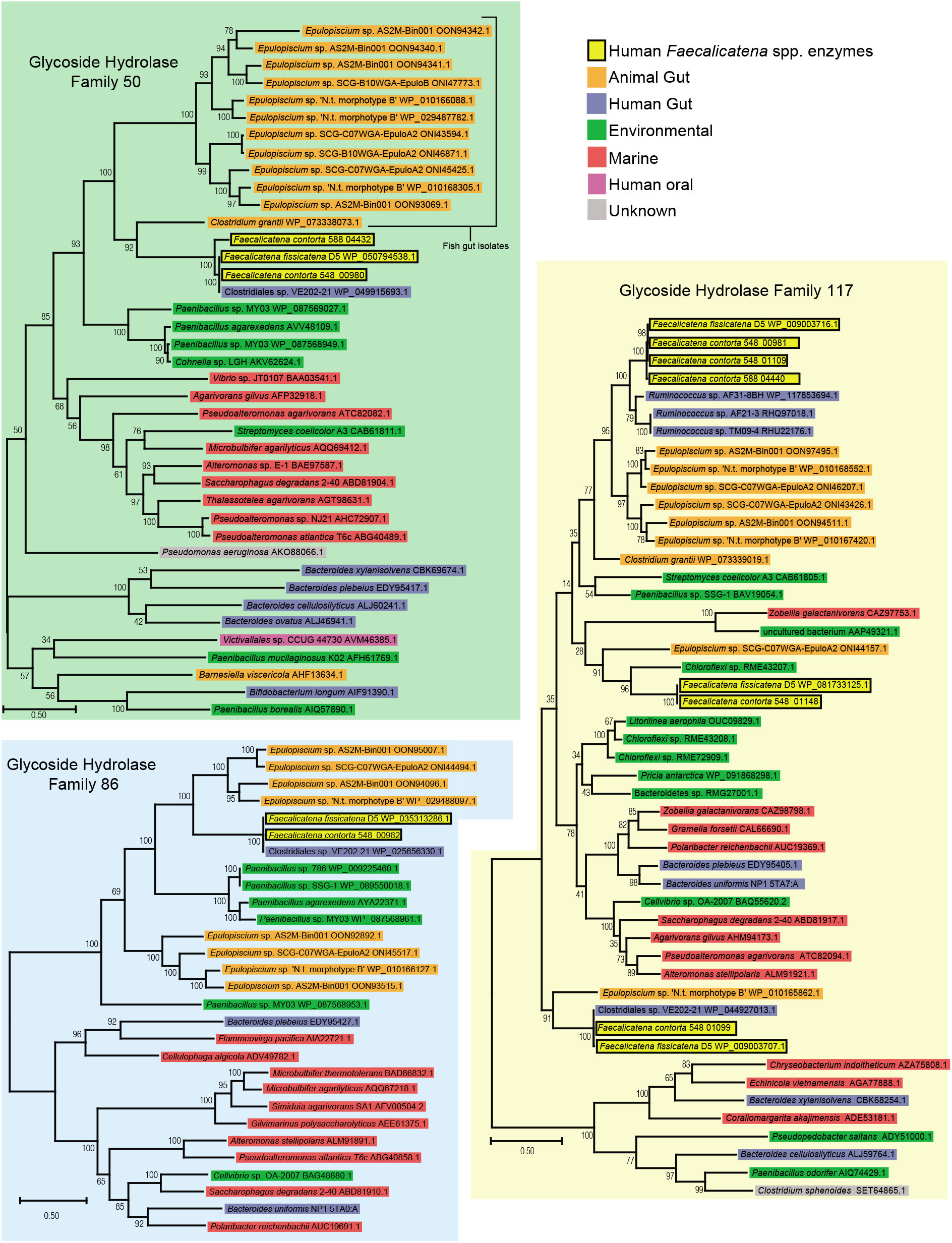
Maximum-Likelihood phylogenic trees generated using deduced amino acid sequences of the GH50, GH86 and GH117 glycoside hydrolases from *Faecalicatena* spp. as well as similar sequences located by BLAST or present in the CAZY database and Protein Data Bank (PDB). Note that the *Faecalicatena* spp. enzymes are often in clades containing enzymes from the marine fish gut symbiont genera *Epulopiscium* and also *Paenibacillus* spp. environmental bacteria that have been isolated from marine sediments. Since the *Clostridiales* sp. VE202-21 enzymes clustered close to the *Faecalicatena* spp. enzymes, we searched public databases to find that this is also a *Faecelicatena contorta* strain. Three human gut *Ruminococcus* sp. enzymes also clustered closely to one of the GH117 enzymes from *Faecalicatena* spp. However, examination of these genomes provided little evidence for similar seaweed degrading phenotypes based on the surrounding genetic architecture.

## Extended Data Table Legends

**Extended Data Table 1.** Seaweed growth and rate data for 354 isolates screened during a high throughput enrichment survey. Strains profiled in this study.

**Extended Data Table 2.** RNA-seq analysis of *Bt*^3731^ genes with a ≥10-fold differential expression when grown on galactose and carrageenan.

**Extended Data Table 3.** BLAST results of the genes in the *Bt*^3731^ CGN PUL showing the closest relative species, their isolation associations and Gram stain reactions.

**Extended Data Table 4a.** Proteomics analysis of *Bt*^3731^ proteins grown on galactose and carrageenan.

**Extended Data Table 4b.** Concordance of proteomic and RNA-seq analysis of *Bt*^3731^ annotated gene products grown on galactose and carrageenan.

**Extended Data Table 5.** RNA-seq analysis of *Bx*^18-002P^ genes with a ≥10-fold differential expression when grown on galactose and porphyran.

**Extended Data Table 6.** BLAST results of the genes in the *Bx*^18-002P^ POR PUL showing the closest relative species, their isolation associations and Gram stain reactions.

**Extended Data Table 7.** Predicted activity of glycoside hydrolases using the SACCHARIS pipeline analysis.

**Extended Data Table 8.** RNA-seq analysis of *Fc*^548^ genes with a ≥10-fold differential expression when grown on galactose, agarose and porphyran.

**Extended Data Table 9.** RNA-seq analysis of *Fc*^588^ genes with a ≥10-fold differential expression when grown on galactose and porphyran.

**Extended Data Table 10.** RNA-seq analysis of *Ff*^D5^ genes with a ≥10-fold differential expression when grown on galactose, agarose and porphyran.

**Extended Data Table 11.** BLAST results of the genes in the *Fc*^548^ AGAR/POR loci showing the closest relative species, their isolation associations and Gram stain reactions.

**Extended Data Table 12.** BLAST results of the genes in the *Fc*^588^ POR loci showing the closest relative species, their isolation associations and Gram stain reactions.

**Extended Data Table 13.** BLAST results of the genes in the *Ff*^D5^ AGAR/POR loci showing the closest relative species, their isolation associations and Gram stain reactions.

**Extended Data Table 14.** Growth media used to culture the isolates in this study.

## References

1 Hehemann, J. H., Boraston, A. B. & Czjzek, M. A sweet new wave: structures and mechanisms of enzymes that digest polysaccharides from marine algae. Curr Opin Struct Biol 28, 77–86, doi:10.1016/j.sbi.2014.07.009 (2014).

2 Porter, N. T. & Martens, E. C. The Critical Roles of Polysaccharides in Gut Microbial Ecology and Physiology. Annual review of microbiology 71, 349–369, doi:10.1146/annurev-micro-102215-095316 (2017).

3 Hehemann, J. H., Kelly, A. G., Pudlo, N. A., Martens, E. C. & Boraston, A. B. Bacteria of the human gut microbiome catabolize red seaweed glycans with carbohydrate-active enzyme updates from extrinsic microbes. Proceedings of the National Academy of Sciences of the United States of America 109, 19786–19791, doi:10.1073/pnas.1211002109 (2012).

4 Shepherd, E. S., DeLoache, W. C., Pruss, K. M., Whitaker, W. R. & Sonnenburg, J. L. An exclusive metabolic niche enables strain engraftment in the gut microbiota. Nature 557, 434–438, doi:10.1038/s41586-018-0092-4 (2018).

5 Pluvinage, B. et al. Molecular basis of an agarose metabolic pathway acquired by a human intestinal symbiont. Nature communications 9, 1043, doi:10.1038/s41467-018-03366-x (2018).

6 Thomas, F. et al. Characterization of the first alginolytic operons in a marine bacterium: from their emergence in marine Flavobacteriia to their independent transfers to marine Proteobacteria and human gut Bacteroides. Environ Microbiol 14, 2379–2394, doi:10.1111/j.1462-2920.2012.02751.x (2012).

7 Mathieu, S. et al. Ancient acquisition of “alginate utilization loci” by human gut microbiota. Scientific reports 8, 8075, doi:10.1038/s41598-018-26104-1 (2018).

8 Déjean, G. et al. Synergy between cell-surface glycosidases and glycan-binding proteins dictates the utilization of specific beta(1,3)-glucans by human gut Bacteroides. mBio (2020).

9 Grondin, J. M., Tamura, K., Dejean, G., Abbott, D. W. & Brumer, H. Polysaccharide Utilization Loci: Fueling Microbial Communities. J Bacteriol 199, doi:10.1128/JB.00860-16 (2017).

10 Hehemann, J. H. et al. Transfer of carbohydrate-active enzymes from marine bacteria to Japanese gut microbiota. Nature 464, 908–912, doi:10.1038/nature08937 (2010).

11 Loureiro RR, Cornish ML & C, N. Applications of carrageenan: with special reference to iota and kappa forms as derived from the Eucheumatoid seaweeds. (2017).

12 Ficko-Blean, E. et al. Carrageenan catabolism is encoded by a complex regulon in marine heterotrophic bacteria. Nature communications 8, 1685, doi:10.1038/s41467-017-01832-6 (2017).

13 Onderdonk, A. B. The carrageenan model for experimental ulcerative colitis. Prog Clin Biol Res 186, 237–245 (1985).

14 Tobacman, J. K. Review of harmful gastrointestinal effects of carrageenan in animal experiments. Environ Health Perspect 109, 983–994, doi:10.1289/ehp.01109983 (2001).

15 Martens, E. C., Kelly, A. G., Tauzin, A. S. & Brumer, H. The devil lies in the details: how variations in polysaccharide fine-structure impact the physiology and evolution of gut microbes. Journal of molecular biology 426, 3851–3865, doi:10.1016/j.jmb.2014.06.022 (2014).

16 Cuskin, F. et al. Human gut Bacteroidetes can utilize yeast mannan through a selfish mechanism. Nature 517, 165–169, doi:10.1038/nature13995 (2015).

17 Zhong, Z. et al. Sequence analysis of a 101-kilobase plasmid required for agar degradation by a Microscilla isolate. Appl Environ Microbiol 67, 5771–5779, doi:10.1128/AEM.67.12.5771-5779.2001 (2001).

18 Poyet, M. et al. A library of human gut bacterial isolates paired with longitudinal multiomics data enables mechanistic microbiome research. Nat Med 25, 1442–1452, doi:10.1038/s41591-019-0559-3 (2019).

19 Jones, D. R. et al. SACCHARIS: an automated pipeline to streamline discovery of carbohydrate active enzyme activities within polyspecific families and de novo sequence datasets. Biotechnol Biofuels 11, 27, doi:10.1186/s13068-018-1027-x (2018).

20 Viana, A. G. et al. beta-D-(1-->4), beta-D-(1-->3) ‘mixed linkage’ xylans from red seaweeds of the order Nemaliales and Palmariales. Carbohydrate research 346, 1023–1028, doi:10.1016/j.carres.2011.03.013 (2011).

21 Arata, P. X., Quintana, I., Raffo, M. P. & Ciancia, M. Novel sulfated xylogalactoarabinans from green seaweed Cladophora falklandica: Chemical structure and action on the fibrin network. Carbohydrate polymers 154, 139–150, doi:10.1016/j.carbpol.2016.07.088 (2016).

22 Browne, H. P. et al. Culturing of ‘unculturable’ human microbiota reveals novel taxa and extensive sporulation. Nature 533, 543–546, doi:10.1038/nature17645 (2016).

23 Hehemann, J. H., Smyth, L., Yadav, A., Vocadlo, D. J. & Boraston, A. B. Analysis of keystone enzyme in Agar hydrolysis provides insight into the degradation (of a polysaccharide from) red seaweeds. J Biol Chem 287, 13985–13995, doi:10.1074/jbc.M112.345645 (2012).

24 Sugano, Y., Kodama, H., Terada, I., Yamazaki, Y. & Noma, M. Purification and characterization of a novel enzyme, alpha-neoagarooligosaccharide hydrolase (alpha-NAOS hydrolase), from a marine bacterium, Vibrio sp. strain JT0107. J Bacteriol 176, 6812–6818, doi:10.1128/jb.176.22.6812-6818.1994 (1994).

25 Becker, S. et al. Laminarin is a major molecule in the marine carbon cycle. Proceedings of the National Academy of Sciences of the United States of America, doi:10.1073/pnas.1917001117 (2020).

26 Kappelmann, L. et al. Polysaccharide utilization loci of North Sea Flavobacteriia as basis for using SusC/D-protein expression for predicting major phytoplankton glycans. The ISME journal 13, 76–91, doi:10.1038/s41396-018-0242-6 (2019).

27 Ngugi, D. K. et al. Genomic diversification of giant enteric symbionts reflects host dietary lifestyles. Proceedings of the National Academy of Sciences of the United States of America 114, E7592–E7601, doi:10.1073/pnas.1703070114 (2017).

28 Kearney, S. M., Gibbons, S. M., Erdman, S. E. & Alm, E. J. Orthogonal Dietary Niche Enables Reversible Engraftment of a Gut Bacterial Commensal. Cell reports 24, 1842–1851, doi:10.1016/j.celrep.2018.07.032 (2018).

29 Desai, M. S. et al. A Dietary Fiber-Deprived Gut Microbiota Degrades the Colonic Mucus Barrier and Enhances Pathogen Susceptibility. Cell 167, 1339–1353 e1321, doi:10.1016/j.cell.2016.10.043 (2016).

30 Martens, E. C. et al. Recognition and Degradation of Plant Cell Wall Polysaccharides by Two Human Gut Symbionts. Plos Biol 9, doi:10.1371/journal.pbio.1001221 (2011).

31 Martens, E. C., Chiang, H. C. & Gordon, J. I. Mucosal Glycan Foraging Enhances Fitness and Transmission of a Saccharolytic Human Gut Bacterial Symbiont. Cell Host Microbe 4, 447–457, doi:10.1016/j.chom.2008.09.007 (2008).

32 Heinz, E. et al. The genome of the obligate intracellular parasite Trachipleistophora hominis: new insights into microsporidian genome dynamics and reductive evolution. PLoS pathogens 8, e1002979, doi:10.1371/journal.ppat.1002979 (2012).

33 Otto, A., Maass, S., Bonn, F., Buttner, K. & Becher, D. An Easy and Fast Protocol for Affinity Bead-Based Protein Enrichment and Storage of Proteome Samples. Methods Enzymol 585, 1–13, doi:10.1016/bs.mie.2016.09.012 (2017).

34 Reisky, L. et al. Biochemical characterization of an ulvan lyase from the marine flavobacterium Formosa agariphila KMM 3901(T). Appl Microbiol Biotechnol 102, 6987–6996, doi:10.1007/s00253-018-9142-y (2018).

35 Zybailov, B. et al. Statistical analysis of membrane proteome expression changes in Saccharomyces cerevisiae. J Proteome Res 5, 2339–2347, doi:10.1021/pr060161n (2006).

36 Cantarel, B. L. et al. The Carbohydrate-Active EnZymes database (CAZy): an expert resource for Glycogenomics. Nucleic acids research 37, D233–D238, doi:10.1093/nar/gkn663 (2009).

37 Zhang, H. et al. dbCAN2: a meta server for automated carbohydrate-active enzyme annotation. Nucleic Acids Res 46, W95–W101, doi:10.1093/nar/gky418 (2018).

38 Edgar, R. C. MUSCLE: a multiple sequence alignment method with reduced time and space complexity. BMC bioinformatics 5, 113, doi:10.1186/1471-2105-5-113 (2004).

39 Darriba, D., Taboada, G. L., Doallo, R. & Posada, D. ProtTest 3: fast selection of best-fit models of protein evolution. Bioinformatics 27, 1164–1165, doi:10.1093/bioinformatics/btr088 (2011).

40 Price, M. N., Dehal, P. S. & Arkin, A. P. FastTree 2--approximately maximum-likelihood trees for large alignments. PLoS ONE 5, e9490, doi:10.1371/journal.pone.0009490 (2010).

41 Letunic, I. & Bork, P. Interactive Tree Of Life (iTOL) v4: recent updates and new developments. Nucleic Acids Res 47, W256–W259, doi:10.1093/nar/gkz239 (2019).

42 Kozich, J. J., Westcott, S. L., Baxter, N. T., Highlander, S. K. & Schloss, P. D. Development of a dual-index sequencing strategy and curation pipeline for analyzing amplicon sequence data on the MiSeq Illumina sequencing platform. Appl Environ Microbiol 79, 5112–5120, doi:10.1128/AEM.01043-13 (2013).

43 Schloss, P. D. et al. Introducing mothur: open-source, platform-independent, community-supported software for describing and comparing microbial communities. Appl Environ Microbiol 75, 7537–7541, doi:10.1128/AEM.01541-09 (2009).

44 Pruesse, E. et al. SILVA: a comprehensive online resource for quality checked and aligned ribosomal RNA sequence data compatible with ARB. Nucleic Acids Res 35, 7188–7196 (2007).

45 Qin, J. et al. A metagenome-wide association study of gut microbiota in type 2 diabetes. Nature 490, 55–60, doi:10.1038/nature11450 (2012).

46 Yu, J. et al. Metagenomic analysis of faecal microbiome as a tool towards targeted non-invasive biomarkers for colorectal cancer. Gut 66, 70–78, doi:10.1136/gutjnl-2015-309800 (2017).

47 Liu, R. et al. Gut microbiome and serum metabolome alterations in obesity and after weight-loss intervention. Nat Med 23, 859–868, doi:10.1038/nm.4358 (2017).

48 Gu, Y. et al. Analyses of gut microbiota and plasma bile acids enable stratification of patients for antidiabetic treatment. Nature communications 8, 1785, doi:10.1038/s41467-017-01682-2 (2017).

49 He, Q. et al. Two distinct metacommunities characterize the gut microbiota in Crohn’s disease patients. Gigascience 6, 1–11, doi:10.1093/gigascience/gix050 (2017).

50 Zhang, X. et al. The oral and gut microbiomes are perturbed in rheumatoid arthritis and partly normalized after treatment. Nat Med 21, 895–905, doi:10.1038/nm.3914 (2015).

51 Nishijima, S. et al. The gut microbiome of healthy Japanese and its microbial and functional uniqueness. DNA Res 23, 125–133, doi:10.1093/dnares/dsw002 (2016).

52 Lloyd-Price, J. et al. Strains, functions and dynamics in the expanded Human Microbiome Project. Nature 550, 61–66, doi:10.1038/nature23889 (2017).

53 Le Chatelier, E. et al. Richness of human gut microbiome correlates with metabolic markers. Nature 500, 541–546, doi:10.1038/nature12506 (2013).

54 Qin, J. et al. A human gut microbial gene catalogue established by metagenomic sequencing. Nature 464, 59–65 (2010).

55 Smits, S. A. et al. Seasonal cycling in the gut microbiome of the Hadza hunter-gatherers of Tanzania. Science 357, 802–806, doi:10.1126/science.aan4834 (2017).

56 Conteville, L. C., Oliveira-Ferreira, J. & Vicente, A. C. P. Gut Microbiome Biomarkers and Functional Diversity Within an Amazonian Semi-Nomadic Hunter-Gatherer Group. Frontiers in microbiology 10, 1743, doi:10.3389/fmicb.2019.01743 (2019).

57 Boratyn, G. M., Thierry-Mieg, J., Thierry-Mieg, D., Busby, B. & Madden, T. L. Magic-BLAST, an accurate RNA-seq aligner for long and short reads. BMC bioinformatics 20, 405, doi:10.1186/s12859-019-2996-x (2019).

58 Quinlan, A. R. & Hall, I. M. BEDTools: a flexible suite of utilities for comparing genomic features. Bioinformatics 26, 841–842, doi:10.1093/bioinformatics/btq033 (2010).

59 Perez-Riverol, Y. et al. The PRIDE database and related tools and resources in 2019: improving support for quantification data. Nucleic Acids Res 47, D442–D450, doi:10.1093/nar/gky1106 (2019).

60 Larsbrink, J. et al. A discrete genetic locus confers xyloglucan metabolism in select human gut Bacteroidetes. Nature 506, 498–502, doi:10.1038/nature12907 (2014).

